# Structure and RNA-binding of the helically extended Roquin CCCH-type zinc finger

**DOI:** 10.1101/2024.03.01.582905

**Authors:** Jan-Niklas Tants, Lasse Oberstrass, Julia E. Weigand, Andreas Schlundt

## Abstract

Zinc finger (ZnF) domains appear in a pool of structural contexts and despite their small size achieve varying target specificities, covering single-stranded and double-stranded DNA and RNA as well as proteins. Combined with other RNA-binding domains, ZnFs enhance affinity and specificity of RNA-binding proteins (RBPs). The ZnF-containing immunoregulatory RBP Roquin initiates mRNA decay, thereby controlling the adaptive immune system. Its unique ROQ domain shape-specifically recognizes stem-looped cis-elements in mRNA 3’-untranslated regions (UTR). The N-terminus of Roquin contains a RING domain for protein-protein interactions and a ZnF, which was suggested to play an essential role in RNA decay by Roquin. The ZnF domain boundaries, its RNA motif preference and its interplay with the ROQ domain have remained elusive, also driven by the lack of high-resolution data of the challenging protein. We provide the solution structure of the Roquin-1 ZnF and use an RBNS-NMR pipeline to show that the ZnF recognizes AU-rich elements (ARE). We systematically refine the contributions of adenines in a poly(U)-background to specific complex formation. With the simultaneous binding of ROQ and ZnF to a natural target transcript of Roquin, our study for the first time suggests how Roquin integrates RNA shape and sequence specificity through the ROQ-ZnF tandem.

## INTRODUCTION

Post-transcriptional gene regulation is an integral part of the cellular processes that control immune responses after pathogen challenge. In particular, the rapid changes in gene expression necessary to activate and differentiate T cells are tightly controlled by RNA-binding proteins (RBPs) governing mRNA stability. Here, Roquin proteins promote target mRNA decay, thereby restricting excessive production of proinflammatory cytokines and preventing inflammatory and autoimmune diseases.

Roquin proteins, Roquin-1 (125.3 kDa) and Roquin-2 (131.3 kDa), are multi-domain proteins that contain a RING domain, a unique ROQ domain, and a zinc finger (ZnF) in their N-terminal half, whereas the C-terminal half is largely unstructured. The CCCH-type ZnF has been identified by sequence analyses and made name-giving for the respective genes, *Rc3h1* and *Rc3h2*. While the Roquin N-terminus mediates target recognition by binding to the 3’-untranslated regions (UTRs) of mRNAs, the C-terminus interacts with the mRNA decay machinery, initiating mRNA deadenylation and decapping upon RNA binding. In the immune system, Roquin proteins control the expression of major regulators of T cell function, such as *ICOS*, *Ox40*, *TNF* and *NFKBID*(*1*). However, Roquin proteins are ubiquitously expressed. Thus, in addition to immune-specific mRNAs, they recognize a plethora of target mRNAs, suggestive of a still little explored housekeeping function in mRNA decay(2).

One outstanding question in understanding post-transcriptional gene regulation is how RBPs recognize their targets with sufficiently high specificity and affinity within the excessive pool of cellular transcripts, to allow for a fine-tuned, cell-type specific mRNA control. Here, precise knowledge of RNA binding preferences is critical to discern direct mRNA targets. With their ROQ domain, Roquin proteins recognize the shape of small RNA hairpins, the so-called constitutive (CDEs) (3,4) or alternative decay elements (ADEs)(2,5,6). Both, ADEs and CDEs, are recognized with low to high nanomolar affinity and are indicators of Roquin targets(6). In contrast to the ROQ domain, the contribution of the ZnF to RNA recognition and regulation is not precisely understood (**Figure 1A**). While it is dispensable for the regulation of *Nfκbiz*(7), it contributes to the recognition of the *A20*(2). Previous ITC experiments with short U-rich RNA oligonucleotides show a low affine binding of ∼35 µM(7). Accordingly, the RNA binding preferences of the Roquin ZnF are currently unclear and thus, our understanding of Roquin-RNA recognition is incomplete.

**Figure 1.**
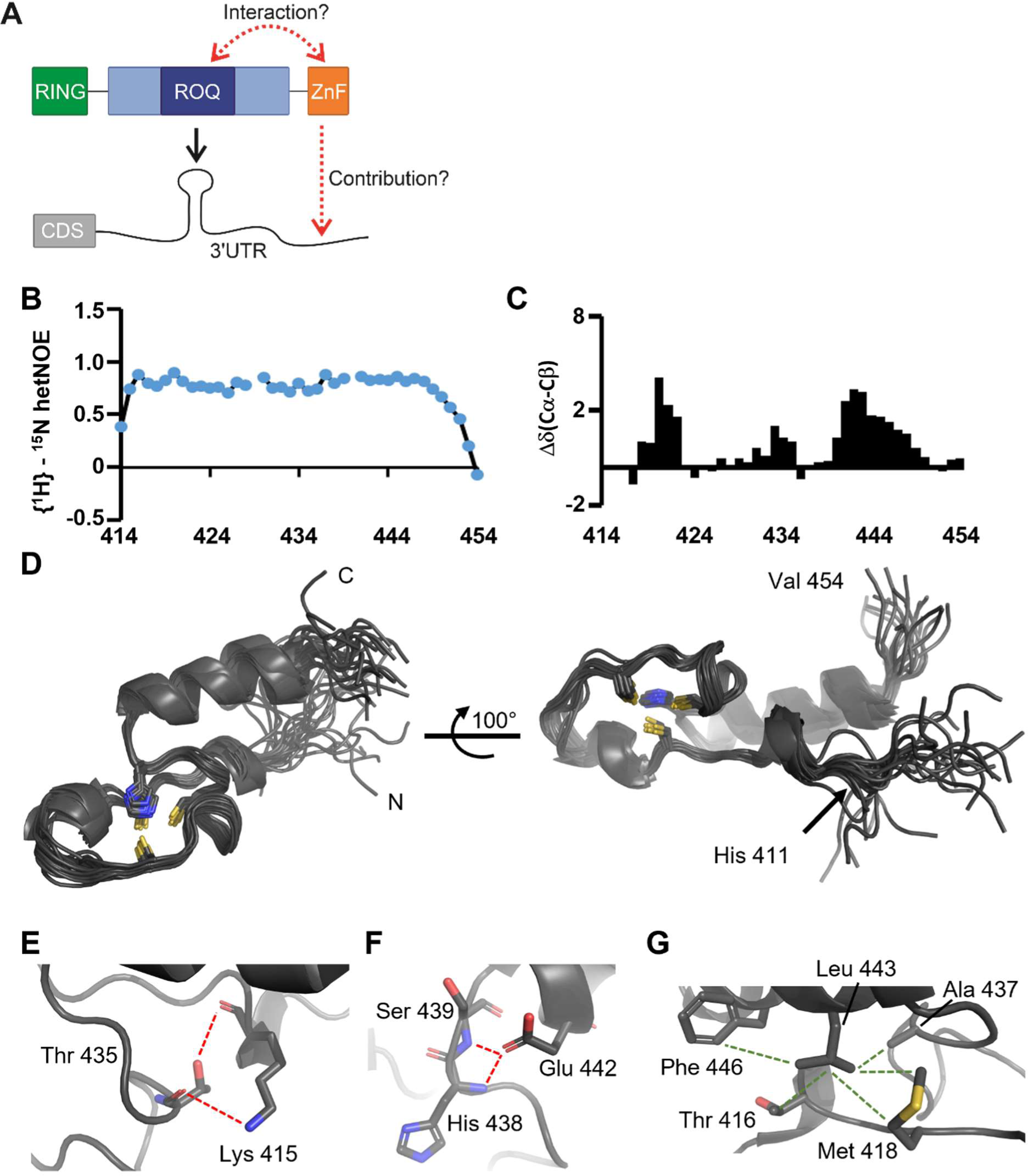
Structure of Roquin-1 CCCH-type ZnF. **A)** Modular architecture of Roquin N-terminus comprising the RING, ROQ and ZnF domains. Dark and light blue show the core and extended ROQ domain, containing the A and B-site required for stem-loop and dsRNA binding in the 3’UTRs of target mRNAs. Domain interactions within Roquin as well as ZnF contributions to RNA recognition are unknown. **B)** {^1^H}-^15^N hetNOE plot of the ZnF spanning residues 414-454. Gaps are derived from proline or non-analyzable amides. **C)** Carbon secondary chemical shifts of ZnF residues 413-454. **D)** Depiction of the 20 lowest-energy structural models of the human Roquin-1 ZnF domain (411-454) including four N-terminal artificial residues. The first natural residue’s position is shown in the right view. The zinc coordinating cysteines 419, 428 and 434 and histidine 438 are shown as sticks. **E-G)** Zoom-in views depicting contacts that stabilize the ZnF fold via polar interactions between K415 and T435 (E), E442 with S439/H438 (F), and a hydrophobic cluster centered on L443 (G).

ZnF domains have become popular for their ability to recognize dsDNA motifs in endonucleases, later on used as tool in modern lab work(8–10). However, ZnF domains can interact with nucleic acids or proteins. While the CCHC and CCHH ZnFs are mostly used to direct protein-protein interactions(11,12), CCCH-type ZnFs facilitate nucleic acid recognition. Typically, CCCH ZnFs prefer unstructured RNAs, often with a preference for AU-rich sequences. As a consequence, many ZnFs have been found to bind AU-rich elements (AREs) in mRNAs, which are known sequence elements to control mRNA decay(13,14). Based on knowledge of high-resolution structures showing complexes of CCCH ZnFs with ssRNAs, there is no obvious hindrance for these representatives to also interact with ssDNA, albeit with a different affinity and recognition mode. However, it may be assumed that distinct binding of DNA or RNA is subordinated to the respective cellular localization of the proteins.

Tight binding of zinc by ZnFs is crucial as it provides the necessary fold that positions side chains for specific interactions. Consequently, the loss of zinc usually is accompanied with functional impairment, as can easily be seen in *in vitro* experiments(15,16).

Interestingly, many ZnFs were found to be challenging in recombinant expression, solubility and stability, which has hindered them from structure determination and systematic analysis of interactions(17). The small folds are often little structured but flexible, while at the same time stable zinc binding is difficult to achieve *in vitro* from recombinant systems. As a consequence, their analysis has often been carried out indirectly in the full-length context, as it was done for e.g. the Roquin ZnF(2). This approach, however, beares the danger of underestimating the precise requirements of ZnFs e.g., because the contribution to a combined RNA motif may appear hidden behind the dominance of more affine RNA-binding domains (RBDs). Attempts to purify an isolated Roquin ZnF in a soluble form and its crystallization within the Roquin N-terminus have failed. While an earlier study has successfully used the ZnF in NMR and ITC experiments, no residue- or atom-resolved information could be provided(7). In the past, all Roquin domains (i.e. the RING domain, a C-terminal part (18)), and the core and extended ROQ domain) have been resolved at high resolution either in separated apo forms (4,19), with RNAs(4,5,20,21), or even as combined domains (PDB entry 4TXA(22)). Still, we lack information of the ZnF fold and its precise interaction sites with potential target RNA. Similarly, we currently have no insight into Roquin’s domain-domain interactions including the ZnF.

The precise definition of domain boundaries and a robust quantification of the functionally relevant fold requires *in vitro* experiments, which also allow to systematically analyze interactions with ligands and the role of RNA sequence. NMR spectroscopy is perfectly suited to investigate proteins with sizes of ZnFs(23–26), e.g. it allows the residue-resolved analysis of a ZnF-RNA complex. In an unbiased approach, the initial lead motif can be obtained in a massive sequencing approach after enrichment from an RNA input pool by the RBD. One such method is RNA-Bind-n-Seq (RBNS), which we exploit as experimental step upstream of a subsequent NMR analysis.

For the first time, we here provide the solution structure of the Roquin CCCH ZnF. We find that the domain contains the expected CCCH zinc coordination, extended by a stabilizing C-terminal helix. We show preferential binding to AU-rich RNA, based on a highly enriched 6-mer derived from RBNS. We find a comparative RNA- and DNA-binding capacity and mode of interaction, which enables a principal component analysis to reveal the precise RNA sequence requirements in an NMR-based approach. We prove that RNA binding is dynamic in pure poly(U) stretches, while incorporated adenosines enhance interactions with the ZnF, which favours binding to a single register. We provide evidence for the structural independence of the ROQ and the ZnF domains and finally prove the simultaneous binding of the two domains to a regulatory RNA sequence in the mRNA of the Phenylalanine-tRNA ligase alpha subunit (*FARSA*), a natural target of Roquin. Altogether, this study fills the longstanding gap in missing structural information on the RNA-interacting Roquin N-terminal multi-domain region. It suggests how Roquin integrates RNA shape and sequence specificity for a fine-tuned RNA target search and technically serves a paradigm for the combined strength of RBNS and NMR, applicable to any other elusive RBD.

## MATERIAL AND METHODS

### Protein expression and purification

Murine protein constructs of coreROQ (aa 171-326), extROQ (aa 89-404) and ZnF (aa 411-454) described previously(27) have been used for this study (**Supplementary Table 1, Figure 1A**). The constructs extROQ-ZnF (aa 89-454) and coreROQ-GS-ZnF (aa 171-326_GGGGS_411-454) were generated by PCR and ligation from Roquin N-terminus used previously(27) and purified accordingly. A freshly transformed colony of *E. coli* BL21 (DE3) was used to inoculate an LB pre-culture supplemented with 50 µg/mL Kanamycin and incubated shaking at 37°C overnight. For the zinc-finger proteins 200 µM ZnCl_2_ was added to the medium. 2 mL of the pre-culture were transferred to 1 L of LB or M9 minimal medium containing 1 g/L ^15^N NH_4_Cl with Kanamycin. At an OD_600_ of 0.8, protein expression was induced with 1 mM IPTG and proteins were expressed at 37°C overnight. Cells were harvested and lysed by sonication in 50 mM Tris pH 8.0, 500 mM NaCl, 4 mM β-mercaptoethanol with DNase I and Protease Inhibitor Mix G (SERVA). Purification was done as previously described(27). Briefly, proteins were purified via IMAC (Ni-NTA), TEV-cleaved overnight and purified via a reverse IMAC. The zinc-finger was purified in the presence of 50 µM ZnCl_2_. Proteins were concentrated using Amicon® centrifugal filter units with appropriate molecular weight cutoffs and a final SEC polishing step using a HiLoad S75 16/600 from GE in 1 M NaCl was performed. Proteins were flash frozen, stored at -80°C and buffer exchanged to 150 mM NaCl, 20 mM Tris pH 7.0, 2 mM TCEP and 30 µM ZnCl_2_ prior to use. For RBNS, the ZnF was expressed as a fusion construct with an N-terminal Twin-Strep-tag® (IBA Lifesciences, Göttingen) and purified as the other constructs.

### RBNS assay

RNA for the RBNS input pool was prepared by *in vitro* transcription. As template, a T7 promoter-containing oligonucleotide was annealed to an equimolar quantity of RBNS T7 template oligonucleotide (a random 20-mer flanked by partial Illumina adapters). 500 fmol template were transcribed overnight at 37°C with 200 mM Tris-HCl pH 8.0, 20 mM magnesium acetate, 8% (v/v) DMSO, 20 mM dithiothreitol (DTT), 20 mM spermidine, 4 mM nucleoside triphosphates (NTP) (each) and self-made T7 RNA-polymerase. The RBNS pool was purified by PAGE (polyacrylamide gel electrophoresis). Oligonucleotide sequences are given in **Supplementary Table 2**.

Based on (28) an RBNS assay was performed with the Twin-Strep-tagged ZnF domain and the randomized input RNA pool. First, the protein was equilibrated at three different concentrations (1, 5, 20 µM) in binding buffer (25 mM Tris-HCl pH 7.5, 150 mM KCl, 3 mM MgCl_2_, 3 mM ZnCl_2_, 0.01% Tween, 500 µg/mL BSA, 1 mM DTT) for 30 min at 4°C. Next, the RNA was folded by snap-cooling and added to a final concentration of 1 µM with 40 U Ribonuclease Inhibitor (moloX, Berlin) and incubated for 1 h at room temperature. A pulldown was performed by incubating the RNA/ZnF mixture with 1 μl of washed MagStrep “type3” XT beads (IBA Lifesciences, Göttingen) for 1 h at 4°C. Subsequently unbound RNA was removed by washing three times with wash buffer (25 mM Tris-HCl pH 8.0, 150 mM KCl, 60 μg/ml BSA, 0.5 mM EDTA, 0.01% Tween). Afterwards, the RNA/ZnF complexes were eluted twice with 25 µL of Elution Buffer (wash buffer plus 50 mM biotin). RNA was extracted with the Zymo RNA Clean & extracted RNA was reverse transcribed into cDNA, amplified by PCR to add Illumina adapters (**Supplementary Table 2**) and an index for each concentration (**Supplementary Table 3**), and subjected to deep sequencing (GENEWIZ, Leipzig). NGS data were analyzed according to the RBNS pipeline as described in (29), available at https://bitbucket.org/pfreese/rbns_pipeline/overview. For structural analysis, the average P_paired_ value across the bases of the indicated motif was calculated for each occurrence in the library using *RNAfold*(30). Motifs are then sorted into five bins according to structure probability. The R value is calculated for each bin as the frequency in the pulldown library divided by that in the input library.

### RNAs and DNA oligonucleotides

*FARSA*+/-AU RNAs were produced by *in vitro* transcription using HDV ribozyme for 3’-end homogeneity as described before(27). Briefly, RNAs were transcribed overnight at 37°C with 200 mM Tris-HCl pH 8.0, 20 mM magnesium acetate, 10% (v/v) DMSO, 20 mM dithiothreitol (DTT), 20 mM spermidine, 4 mM nucleoside triphosphates (NTP) (each) and self-made T7 RNA-polymerase and PAGE purified. RNAs were eluted from gels by ‘crush-and-soak’, i.e. gel slices were cut in pieces and incubated rotating at room temperature in 0.3 M sodium acetate pH 6.5. Afterwards, RNAs were precipitated using ethanol/acetone. Finally, the RNA was resolved in double-distilled water and stored at -20°C. For buffer exchange (NMR buffer: 150 mM NaCl, 20 mM Tris pH 7.0, 2 mM TCEP) and concentrating, we used Vivaspin® concentrators (molecular weight cut-off 3 kDa). The RNA was folded by snap-cooling, i.e. after heating to 95°C for 5 min the RNA was stored on ice.

Short RNAs were purchased from Horizon (UAUUAUAU) and Eurofins Genomics (UUUUU, UAUUU, and UAUAU) and dialyzed against NMR (150 mM NaCl, 20 mM Tris pH 7.0, 2 mM TCEP) buffer. All RNAs were heated and snap cooled before use (**Supplementary Table 4**). DNA oligonucleotides used for NMR titrations (**Supplementary Table 4**) were purchased from Sigma Aldrich and dissolved in NMR buffer.

### NMR spectroscopy

All NMR experiments were recorded in the Frankfurt BMRZ and the Bavarian NMR center at TU Munich. For backbone and side chain assignments, we used Bruker Avance spectrometers of 500, 600 and 800 (Avance III) MHz proton Larmor frequencies equipped with triple-resonance cryo-probeheads and operated with TOPSPIN version 3. To this end, we recorded the following experiments using a U-^15^N-labeled sample of 360 µM and a U-^13^C-^15^N-labeled sample of 740 µM: HNCACB; CBCAcoNH, HNCO, HNcaCO, HBHANH, 3D-CCH-TOCSY, 3D-^1^H,^13^C-NOESY-HSQCs (mixing time of 150 ms), 3D-^1^H,^15^N-NOESY-HSQC (mixing time of 135 ms) and a lr-HSQC to identify the Histidine side chain protonation state as described earlier(31).

A ^15^N-heteronuclear NOE experiment was recorded as an HSQC-based version (based on the standard ^1^H,^15^N-HSQC acquisition parameters: nitrogen offset of 117 ppm and a spectral width of 32 ppm, proton spectral width 12 ppm, acquisition time of 84 ms) using a pseudo-3D in an interleaved saturation/no-saturation scanning mode at 600 MHz proton Larmor frequency and including WATERGATE as water suppression(32). The saturation time was set to 3010 ms and 128 complex indirect points were measured for each condition.

All spectra were processed using TOPSPIN 3 versions. Resonance assignments were carried out in CcpNMR Analysis version 2.1 to 2.4(33). Peak picking of all spectra including NOESYs, the quantitative analysis of hetNOEs and secondary chemical shifts were performed with implemented tools of Analysis.

For titration experiments ^1^H,^15^N-HSQC and imino proton spectra of 30 µM apo protein were recorded at 298 K. Additional spectra were recorded in presence of 2.0-fold excess of DNA or 0.5, 1.0 and 1.2-fold amounts of *FARSA*-AU RNA. For the UAUUAUAU RNA, the TATTATAT DNA and the GAGCAC control RNA 0.25, 0.5, 1.0, 1.5, 3.0 and 6.0-fold amounts of nucleic acids were added to the protein. Chemical shift perturbations (CSP) were calculated according to the following formula:

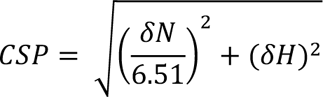

Imino proton resonances of the *FARSA*-AU RNA were assigned with a ^1^H,^1^H-NOESY recorded with a mixing time of 200 sms on a 160 µM sample in NMR buffer at 293 K. 1D proton spectra of *FARSA*-AU RNA variants were recorded at 293 K as well. ^1^H,^1^H-TOCSY experiments were recorded on 200 µM RNA samples complemented with 400 µM ZnF. The mixing time was 30 ms and 80 scans were collected for 400 indirect points at 298 K. All Bruker AV spectrometers used (600, 700 MHz proton Larmor frequencies) were equipped with triple-resonance cryo-probes. For processing, Topspin 4.0.6. (Bruker) was used and spectra were analyzed with NMRFAM-Sparky 1.414(34).

### Structure calculation

For structure calculation (see **Supplementary Table 1**), we used ^1^H-^1^H distance restraints obtained from 3D-^1^H-^13^C-NOESY-HSQCs (individual aliphatic and aromatic, with either a mixing time of 150 ms), 3D-^1^H-^15^N-NOESY-HSQC (mixing time of 135 ms). All NOEs were initially assigned automatically using the CYANA(35) implemented routine. To ensure stable initial folding of the zinc cluster, we included 14 additional restraints between all four coordinating residues and an installed pseudo zinc atom according to the expected standard geometry of CCCH zinc clusters: Cys-Cβ to Zn (3.35-3.45 Å); Cys-S to Zn (2.25-2.35 Å); His-NE2 to Zn (1.95-2.05 Å). No additional restraints were placed between cysteines and the histidine to avoid biases in the 3D cluster geometry. 20 additional distance restraints were included for helicity based on respective ^15^N-NOESY patterns. We also used 57 dihedral angle restraints obtained from TALOS(36). Stereospecific side chain Hβ assignments were defined for residues K415, Y417, M418, C419, C428, C434, H438, and L443.

CYANA v.3.98.13(37) was used to create 100 structures calculated using 32,000 refinement steps per conformer with 20,000 high temperature steps of torsion angle dynamics followed by 20,000 steps of slow cooling. Seven iterative rounds of structural model building and refining were followed by a final round of refinement to assign NOEs and undergo simulated annealing. For the final run the heavy atom RMSD compared to the final ensemble from run 1 to 7 decreased from 2.59 to 0.82 Å. The 20 lowest-energy models were chosen for deposition after removal of the pseudo Zn atoms, while for clarity and to prove convergence they are shown in **Supplementary Figure 1C**. The overall backbone RMSD of the final ensemble for residues 413-451 is 0.64 ± 0.19 Å based on an average of 8.5 restraints per residue. PyMOL (Schrödinger, Inc.) was used for subsequent structure visualization. Further assessment of the structural ensemble has been carried out using the CYANA-created output (no distance and no van der Waals restraints, two minor angular violations, but no Ramachandran outliers) as well as through the PSVS server run by the Montelione lab (only favored and allowed Ramachandran regions, no bond lengths and angle deviations)(38). The final ensemble has been deposited in the PDB (ID: 8rhs) and the BMRB (ID: 52226).

### Sequence alignment

CCCH-type ZnF sequences obtained from the UniProt database(39) were aligned to residues 411-454 of the Roquin ZnF using Dialign(40). The Zn-coordinating amino acids (CCCH) were fixed as anchor points. Flanking residues not aligning with amino acids 411-454 of Roquin were cropped. The conservation score was calculated with SnapGene based on (41,42). The weblogo (consensus motif) was created by the weblogo webserver 2.8.2(43) with the small sample correction option. For multiple species sequence alignment of Roquin-1 Clustal Omega was used(44).

### Isothermal titration calorimetry

All ITC runs were performed on a MicroCal iTC200 (Malvern Instruments) at 20°C. A 500 µM RNA stock (UAUUAUAU) was titrated to 20 µM of protein in the cell. The stirring speed was 750 rpm, the initial delay was 120 s and the spacing time between injections 120 s. The reference power was set to 11 µcal^-1^. Injection volumes were 0.2, 5 x 0.5 and 20 x 2 µl; in total 26 injections. Protein and RNA were dialyzed in the NMR buffer overnight prior to measurement. Data were processed with NITPIC 1.2.7(45,46). Curve fitting was done in SEDPHAT(47) using ΔH values obtained from NITPIC. Values shown in **Figure 2G** correspond to averages and standard deviations from n = 3.

**Figure 2.**
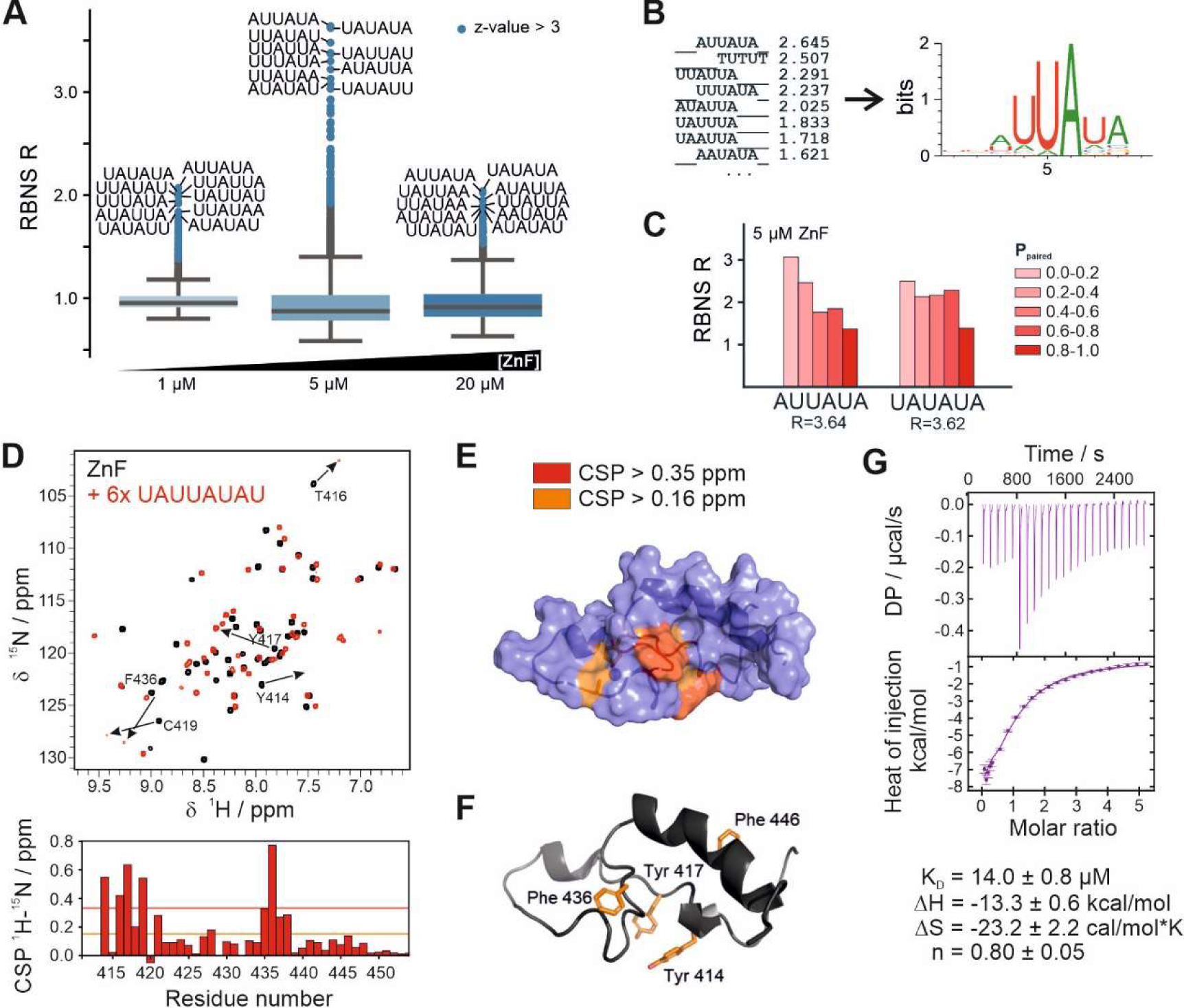
RBNS identifies single-stranded, AU-rich sequences as binding motifs for the ZnF of Roquin-1. **A)** Enrichment of all 6-mers at 1, 5 and 20 µM ZnF concentration. Values greater than three standard deviations above the mean are highlighted. For the highest ten motif sequences are given. **B)** Composite logo containing the significantly enriched 6-mers. **C)** Structural features of the two most enriched motifs at 5 µM protein Motifs are sorted into five bins according to structure probability. **D)** ^1^H,^15^N-HSQC spectra of apo ZnF (black) and in complex with AU-rich RNA (red). Below calculated CSPs are plotted against residue number. Orange and red lines indicate the average plus one and two standard deviations, respectively. **E)** CSPs mapped onto ZnF structure. CSPs larger than two (red) and one (orange) standard deviation are visualized. **F)** Steric distribution of aromates in the Roquin-1 ZnF domain. The protein is shown as cartoon with conserved aromatic residues shown as sticks in orange. The zinc-coordinating His 438 and the unstructured His 411 are left out. **G)** ITC curve of AU-rich RNA binding by ZnF.

### Electrophoretic mobility shift assays (EMSAs)

RNAs were dephosphorylated using Quick CIP (NEB) at 37°C. After 5’-end-labeling with [γ-^32^P]-ATP (Hartmann Analytics) and T4 PNK, RNAs were passed through NucAway Spin columns (Invitrogen) for purification. RNAs were refolded in NMR buffer by snap cooling and stored at -20°C until usage. For EMSAs, 2 µl of target RNA were incubated for 15 min with 0.6 µg baker’s yeast tRNA_Phe_ (Roche), 1 mM MgCl_2_ and increasing amounts of protein in a volume of 20 µl. Protein concentrations of 0, 20, 50, 100, 150, 200, 300, 400, 500, 700, 1.000, 2.000, 5.000, 10.000, 20.000 nM were used. Samples were mixed with loading buffer (1x TB, 60% glycerol, 0.02% Bromphenolbue) and run on 6% polyacrylamide gels as described before(27). Gels were dried, exposed to a phosphor storage screen and imaged on a Typhoon 9400 Variable Mode Imager (GE). For the quantitative EMSA of extROQ-ZnF vs. *FARSA*-AU RNA, 5 µM of unlabeled *FARSA*-AU RNA was spiked with 3 nM ^32^P-labeled *FARSA*-AU RNA and incubated with increasing amounts of protein (0, 1, 2, 3, 5, 7.5, 10, 15 µM). The decrease of free RNA was quantified and plotted as described before(27).

## RESULTS

### Structure of Roquin ZnF

Earlier work had failed to determine the atomic structure of the Roquin ZnF domain. Initial studies had been based on the predicted domain boundaries between natural residues 411 to approximately 440, likely guided by standard algorithms centered on the CCCH pattern. A respective construct turned out completely insoluble in our hands (data not shown), while a Roquin 1-440 construct appeared soluble and sufficiently stable when expressed from *E.coli*. In 2018(7), we used a C-terminally extended version of the ZnF domain (411-454) and found this protein version stably expressing in a soluble manner by help of a solubilizing thioredoxin tag in additional presence of zinc. The purified sample revealed a folded moiety judged by NMR-based observations. We decided to use this construct for *en-detail* NMR-based analyses and RNA-binding studies with residue resolution.

To this end, we set out to assign NMR backbone resonances of a ZnF construct carrying an N-terminal GAMA-overhang derived from an included TEV cleavage site. To our surprise, we were unable to unambiguously assign amide groups to residues before Y414, i.e. we faced intense line-broadening for resides G(-4)-K413 indicating conformational averaging, increased flexibility (lack of structure), or exchange with water. This is supported by a hetNOE plot shown in **Figure 1B**, where we still find residue Y414 significantly less structured then the rest of the domain including residue 450. Apart from this, we were able to achieve complete backbone assignment of residues 414-454. To get a first impression of the underlying structural context, we used secondary carbon chemical shift analysis for residues 413-454 (**Figure 1C**). Unexpectedly, we found a prolonged helical C-terminal extension (440-451) after the CCCH zinc cluster (expected to be formed by C419, C428, C434, and H438), which indicates a crucial secondary structural element is needed for the stable ZnF composition, and likely solubility.

Driven by this observation, we decided to determine the solution structure of the ZnF and first carried out the necessary side chain assignments. Similar to the observations for the backbone, we were only able to assign approximately 85% of all possible atoms (including the backbone), but achieved more than 90% for the region 413-454, indicating that some of the more structured regions still undergo substantial dynamics that renders respective side chain resonance invisible. Those numbers, however, were sufficient to determine the structure that shows a good convergence of the 20 lowest-energy ensemble members for region 413-451 (**Figure 1D, Supplementary Figure 1C,** and **Table 1**). In line with our prior observation, the rigid domain starts from residue K413. The structure is centered on the CCCH zinc-coordinating cluster (see also **Supplementary Figure 1A**), followed by a canonical 12-residue α-helix with decreasing convergence, in line with the SCS plot in **Figure 1C** that indicates decreasing helicity to the C-terminus.

**Table 1:**
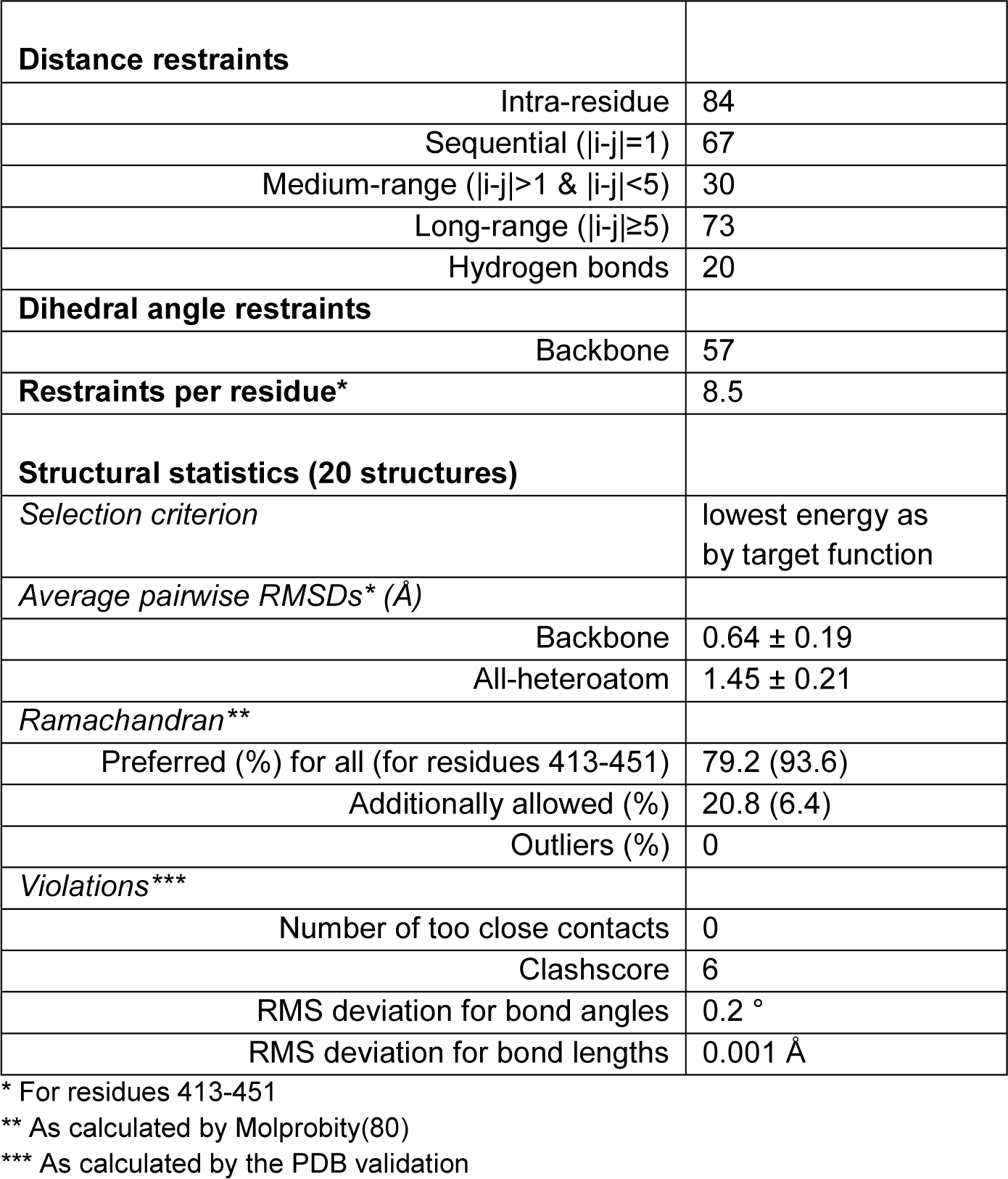
Structural statistics of Roquin-1 ZnF NMR structure.

The zinc-coordinating cluster is embedded into the domain fold through only few contacts, e.g. with the highly conserved K415 interacting with T435 (**Figure 1E**). Additionally, the side chain of E442 stabilizes the cluster with the N-terminal helical end (**Figure 1F**). Interestingly, the helix shows only little stabilizing contacts with the N-terminal part of the ZnF fold, mainly mediated by van der Waals and hydrophobic interactions around L443 (**Figure 1G**). Nonetheless, we suggest this very helical extension has escaped prior predictions, but appears as an essential and unique element of the Roquin-1 ZnF, and consequently this region is highly conserved between species (**Supplementary Figure 1B**). Remarkably, in the process of structure determination, we found that AlphaFold(48) was in fact capable of predicting the helix in an almost identical geometric context of the Roquin ZnF overall fold (see comparison in **Supplementary Figure 1D**). Of note, AlphaFold also was able to predict a short helical element within the CCCH zinc cluster region, which likely serves to restraint the zinc-engaging loop on one side and thus allowing for a stably coordinated metal and a robust domain. While AlphaFold precisely matches the predicted fold, we find minor deviations for the N-terminal end and the concrete zinc binding moiety (no metal is included in AlphaFold though). Altogether, we here for the first time provide the atomic structure of the last unexplored Roquin domain and thus complement the so-far missing piece to the overall architecture and ultra-conserved region (**Supplementary Figure 1B**) of the folded Roquin N-terminal half, responsible for RNA-recognition.

### The ZnF recognizes AU-rich RNA motifs

We used RNA Bind-n-Seq (RBNS(28)) to systematically explore the binding specificity of the ZnF. This *in vitro* high-throughput assay provides the ability to identify the recognized sequence motifs by an RBP, potentially followed by a secondary structure prediction (**Supplementary Figure 2A**). Thus, a 20-nt random RNA pool flanked by short constant adapter sequences was incubated with different concentrations of the Strep-tagged ZnF domain (1, 5, 20 µM). NMR spectroscopy was used to confirm structural integrity of the tagged fusion protein (**Supplementary Figure 2D**). The constant sequence regions were used to add sequencing adapters for subsequent analysis of bound RNAs by Next Generation Sequencing.

From sequencing, we obtained ∼40-50 million unique reads for each protein concentration. By comparing with a similar number of reads from the library of the input RNA pool, we identified enriched k-mers of six nucleotides (k=6) for each concentration. We calculated enrichment values (“R”), which are defined as the frequency of a k-mer in the reads of the pulldown divided by its frequency in the input reads (**Figure 2A, Supplementary Table 5**). Analysis of enriched 4-, 5- and 7-mers yielded similar results **(Supplementary Figure 2B and Supplementary Table 5)**. The most enriched motifs are characterized by alternating adenines and uracils, where the distance between two adenines is one or two uracils. The most enriched 6-mers cluster into a single AU-rich motif with a central UA dinucleotide (**Figure 2B**). Structural features of the identified binding motifs were predicted by calculating the average base pairing probability. For the most enriched motifs, the highest R-value is found at lower folding probabilities (**Figure 2C, Supplementary Figure 2C**). Thus, the RBNS assay identifies AU-pure, single-stranded regions as preferred binding motifs for the ZnF domain of Roquin-1. However, we are still lacking the precise sequence requirements of the domain.

### Binding mode of AU-rich RNA

To confirm and in-depth characterize the ZnF interaction with AU-rich RNAs on a molecular level, we used the RBNS-derived sequence AUUAUA for *in vitro* experiments. Based on typical ZnF motif lengths of 3-4 nts (49–51), our RBNS motif potentially allows multiple target registers. Despite, and as we do not know the precise motif length required, we extended the RNA with two flanking uracils, thereby providing additional motif options. For residue-resolved information, we recorded ^1^H,^15^N-HSQC NMR spectra of RNA-protein complexes. The observed major chemical shift perturbations (CSP, **Figure 2D**, **Supplementary Figure 2E**) in presence of the UAUUAUAU RNA indicate binding of the ZnF. Quantification of CSPs reveals key residues for interaction, which -as expected-cluster in proximity to the zinc coordination site (**Figure 2E**). In particular, the aromatic residues Y414, Y417 and F436 exhibit large CSPs (>0.35 ppm). Both amino acids preferentially interact with uracils and adenines, respectively, through base-stacking (52). We suspect the C-terminal helix to sterically shield F446 from interacting with RNA (**Figure 2F**). Of note, we do not observe significant CSPs for this helix upon RNA addition. Nevertheless we cannot rule out that it contributes to RNA binding due to its charges (rich in R, L at the C-terminus) in a full-length protein context. A control RNA devoid of uracils (GAGCAC) only leads to smaller changes in the spectra, supporting the ZnF preference for AU-rich sequence motifs (**Supplementary Figure 2F and G**). Careful inspection of the UAUUAUAU titration series (**Supplementary Figure 2E**) unveils that most ZnF peaks shift in the fast exchange regime in the presence of RNA. In addition, we observed line-broadening for a few residues at substoichiometric RNA amounts. This fast-to-intermediate exchange regime suggests a low-micromolar affinity of the ZnF for AU-rich RNAs. We used isothermal titration calorimetry (ITC, **Figure 2G**) to obtain a precise affinity and indeed confirmed the NMR-based low-micromolar interaction (K_D_ = 14.0 ± 0.8 µM) and a 1:1 stoichiometry. This is in agreement with previously described affinities of RNA binding CCCH-type ZnFs (53) and confirms the Roquin ZnF as a specific single-strand AU-rich RNA binder.

### Mode of RNA and DNA binding

We next set out to understanding nucleic acid sequence composition required for Roquin ZnF interaction. As crystallization trials failed and the moderate affinity of the RBNS motif impedes an unambiguous NMR structural analysis of complexes, we tested a variety of sequences by NMR. ZnFs usually bind exclusively DNA or RNA, and little is known about dual recognition(54) or equal binding of DNA and RNA(55). We asked whether the Roquin ZnF binds AT-rich DNAs(56). Hence, we first tested an RNA-analogous TATTATAT DNA oligonucleotide for ZnF binding and compared it to the respective RNA. The NMR titration (**Figure 3A** and **Supplementary Figure 3A**) shows chemical shift differences indicative of binding. CSP plots reveal that DNA binding occurs through the same interface as RNA binding, as the same residues exhibit strong chemical shift changes in presence of the nucleic acid (compare **Figure 2D** and **Supplementary Figure 3B**). Both data sets show a good correlation, which supports a common binding interface (**Supplementary Figure 3D**). Interestingly, some ZnF residues respond differently to RNA or DNA, i.e. their trajectories or the magnitudes of shifts differ (**Figure 3B** and **Supplementary Figure 3C**). These residues are therefore potentially sensitive for 2’-OH groups. Of note, in presence of DNA line broadening is neglectable, compared to the RNA titration (**Supplementary Figure 3A**). DNA binding occurs thus in pure fast exchange which suggests weaker binding of the DNA oligonucleotide. We conclude that the Roquin ZnF prefers RNA over DNA, but it can bind both nucleic acids through a common mode (**Supplementary Figure 3D**). This feature can be exploited in binding studies, where the use of DNA oligonucleotides offers advantages over RNA: DNA oligonucleotides can be obtained commercially in relevant quantities compared to the laborious in-house production of RNA. Together with the low costs of DNA oligonucleotides this allows to test more sequences.

**Figure 3.**
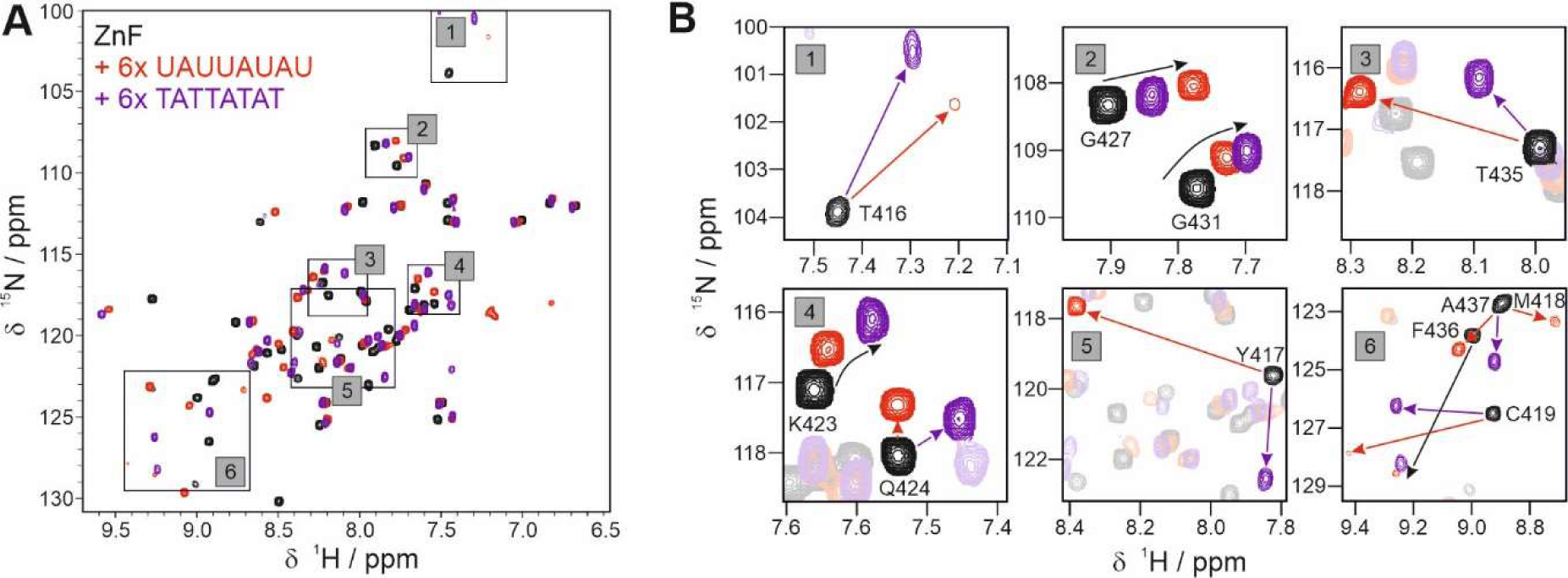
Key ZnF residues involved in RNA and DNA binding. **A)** ^1^H,^15^N-HSQC spectrum of apo ZnF (black), and in presence of RNA (red) or DNA (purple). **B)** Zoom-ins of A) highlighting individual key residues. Arrows indicate CSPs upon addition of the respective nucleic acid.

### Contribution of adenines to ZnF binding

We aimed to narrow down our RBNS-derived consensus motif to dissect individual contributions of nucleobases for RNP formation, i.e. frequency and arrangement of uracils and adenins. By NMR spectroscopy, we analysed ZnF chemical shift perturbations in presence of a systematic variation of DNA oligonucleotides (**Figure 4** and **Supplementary Figure 4**). As our RBNS experiments confirmed the ZnF to be an AU-binder, we first tested poly(T) and poly(A) oligonucleotides and poly(C) as a control. While significant changes are observed in the protein spectrum upon addition of poly(T), only marginal differences occur in presence of poly(A) and poly(C) (**Supplementary Figure 4A and B**). We conclude that the ZnF binding site cannot accommodate two neighbouring purines, likely due to steric clashes within the binding pocket. However, the sequence composition of our UAUUAUAU RNA suggests that adenines are bound at specific positions. We therefore tested poly(T) stretches containing one or multiple deoxyadenosines and varied their spacing. Interestingly, for most key residues, we detected larger chemical shift perturbations in DNAs containing one deoxyadenosine (**Figure 4A and C, Supplementary Figure 4**): F436 shows a strong CSP increase in presence of deoxyadenosine (**Figure 4A** and **Supplementary Figure 4C**), which points at stacking interactions. A second deoxyadenosine only leads to a small additional increase, as can be seen by differential CSP plots (**Supplementary Figure 4C**). Here, spacing of the purines is essential: TTTAATTT causes smaller changes than TTATATTT (**Figure 4A and C, Supplementary Figure 4C**), underpinning potential steric hindrance through adjacent purines as suggested above. Larger spacing does not significantly increase CSPs according to our experiments with a TTATTATT and TATTTTAT DNA (**Figure 4C**, **Supplementary Figure 4A and B**). Interestingly, a deoxyadenosine embedded in poly(T) or at the 5’-end causes larger CSPs than a 3’-terminal deoxyadenosine, suggesting preferential binding of the ZnF to -AT-sequences (**Figure 4B and C**, **Supplementary Figure 4A and B**). To our surprise, a single guanine embedded in poly(T) almost abolishes ZnF binding (**Supplementary Figure 4A and B**), which shows its high specificity for AU-tracts. We interpret this reduction in CSPs to mean that incorporation of guanine reduces potential – and putative dynamic – binding registers. Binding of the Roquin ZnF is thus significantly enhanced by single (deoxy)adenosines embedded in poly(T) or poly(U) stretches and requires at least one nucleotide spacing of tandem (deoxy)adenosines.

**Figure 4.**
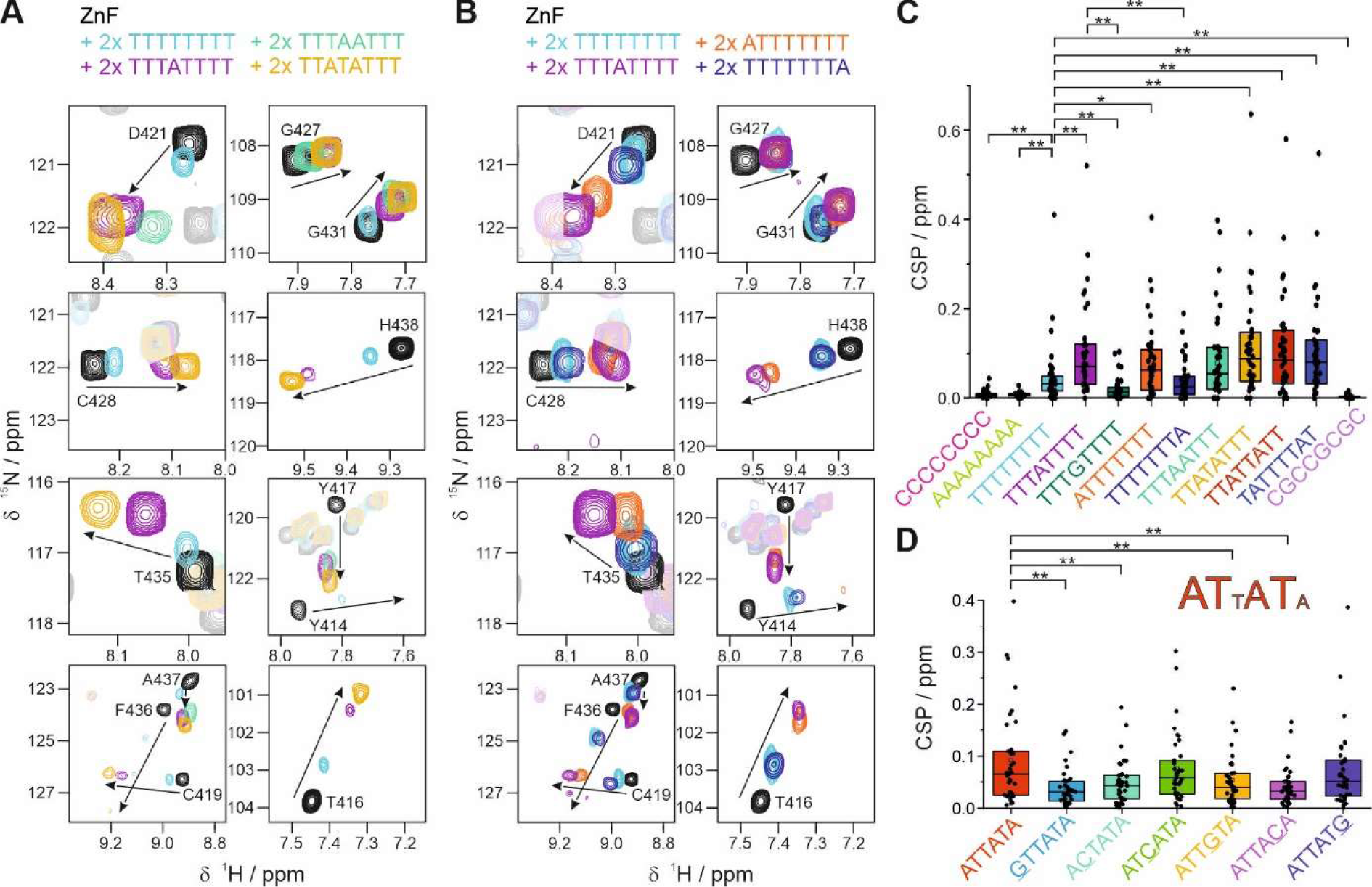
Sequence requirements for nucleic acid binding through Roquin ZnF. **A and B)** Zoom-ins of ^1^H,^15^N-HSQC spectra from Supplementary Figure 4A highlighting individual key residues of apo ZnF (black) and in presence of DNAs listed above (color). Arrows indicate chemical shift perturbations upon addition of the respective DNA oligonucleotide. **A)** shows spectra of ZnF in complex with DNA oligonucleotides with varying A content, while **B)** compares DNA oligonucleotides with varying positioning of As. **C)** Box plot of CSPs calculated from titrations shown in A) and Supplementary Figure 4A and B. Significance of CSP analyses was calculated in a t-test referring to TTTTTTTT or TTTATTTT with significance levels of p<0.01 (**) and p<0.05 (*). **D)** Box plot of CSPs calculated from titrations shown in Supplementary Figure 5A and B. Underlined residues show mutation sites. Font size in the RBNS motif indicates rank and impact on ZnF binding as based on significance level of CSP analysis.

### Relevance of individual positions in the RBNS motif for ZnF binding

Our systematic analysis of a reconstructed AT-rich oligonucleotide does not capture the full complexity of the RBNS-derived motif. We therefore compared single-position mutant versions thereof to the original RBNS-motif in NMR titrations (**Supplementary Figure 5A**). CSP patterns were conserved but appeared smaller for all mutants (**Figure 4D** and **Supplementary Figure 5B**). The mutants at positions 1, 2, 4 and 5 showed a significantly stronger reduction in average CSPs (**Figure 4D**) as well as for the maximum CSPs compared to the RBNS motif and mutant positions 3 and 6. Based on that we ranked the six positions within the RBNS-motif according to their impact on ZnF binding. Interestingly, two AT-dinucleotides have the largest effect on CSPs, with an additional but dispensable thymine in between. This confirms our findings from the bottom-up approach before, that the ZnF prefers 5’ adenines and requires one or two thymines for spacing of multiple adenines.

### Multiple register binding in AREs

Our NMR data indicate that the ZnF binds poly(T)/(U) as well as adenines embedded in poly(T)/(U) stretches comparably well. Further, the length of our model RNA exceeds the number of bases required for ZnF binding(23,49,50). Consequently, we next asked whether binding registers depend on RNA sequence. We made use of ^1^H,^1^H-TOCSY NMR spectra of RNA, which have previously proven valuable for evaluating multiple register binding of a protein(57,58). Each uridine gives rise to a cross-peak based on its H5-H6 coupling. Unstructured poly(U)s will collapse into one single broad peak. However, distinct chemical environments for individual bases, e.g. through protein binding, cause peak dispersion. The number of detectable peaks can thus serve as a readout for stable complex registers. We recorded ^1^H,^1^H-TOCSY spectra on short RNAs (**Figure 5A** and **Supplementary Figure 6**) in presence of a 2-fold excess of ZnF. For UUUUU we observed two broad peaks barely separated from each other, indicating no single complex conformation. This is in line with our observation that guanines perturb complex formation with poly(T) (see above, **Supplementary Figure 4B**), where the beneficial effect of multiple possible registers for complex stability is abolished. In presence of a single (UAUUU) or two (UAUAU) adenines we detected three uracil peaks, which suggests a single binding register (**Figure 5A**). Similarly, for the UAUUAUAU RNA the spectrum shows four signals pointing at both one register and a diversified chemical environment due to the asymmetric sequence. Our NMR data support the specific recognition of a single adenosine within a poly(U) stretch, which leads to a preferred binding register stably locked by the ZnF, e.g. via concerted aromatic stacking (**Figure 5B**).

**Figure 5.**
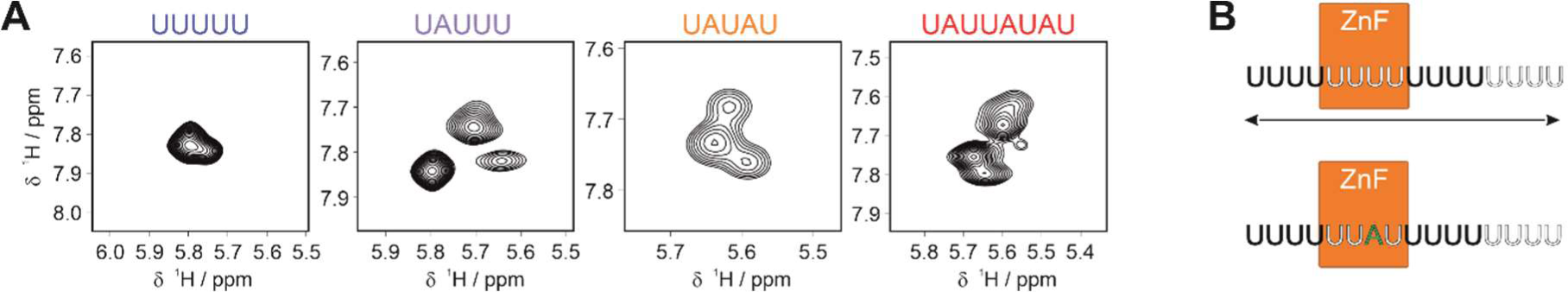
Multiple register binding in poly(U) RNAs. **A)** ^1^H,^1^H-TOCSY sections of RNAs complexed with ZnF showing Uracil H5-H6 correlations. Full spectra are shown in **Supplementary Figure 6A**. **B)** Model of dynamic ZnF binding to poly(U) (top) and single register binding to adenines (bottom).

### Independent RNA-binding by Roquin through the ROQ and ZnF domains

We have shown that the ZnF of Roquin binds AU-rich RNA with low micromolar affinity (**Figure 2G**). Next, we asked, whether RNA binding of the ZnF could be modulated by the neighboring ROQ domain, or *vice versa*. We first assessed whether the domains interact and recorded ^1^H,^15^N correlation NMR spectra (**Supplementary Figure 7B**). Individual domain spectra overlay well with those from the two-domain constructs. Furthermore, addition of extROQ to the ZnF did not lead to alterations in the ZnF fingerprint spectrum. We conclude that the ROQ domain and the ZnF are independent moieties in the Roquin protein apo state. We tested RNA binding with two constructs comprising the two domains. One contained the natural sequence of extROQ-ZnF, the other the coreROQ domain fused with an artificial GS linker to the ZnF to preserve domain independence (**Supplementary Table 1, Supplementary Figure 7A, B**). With these we performed EMSAs to monitor RNA binding through band shifts in native gels.

We used a stem-loop containing fragment from the *FARSA* 3’-UTR (**Supplementary Figure 7C**). *FARSA* encodes for the Phenylalanine-tRNA ligase alpha subunit and provides a CDE flanked by a 5’ AU-rich sequence. It thus combines the two *cis*-elements required for binding of both ROQ and ZnF domain and serves as a model system. Structural integrity of this RNA, which we called *FARSA*-AU, was confirmed by a ^1^H,^1^H-NOESY NMR spectrum (**Supplementary Figure 7C**), where the presence of imino proton peaks confirmed formation of base-pairs as predicted. The isolated domains extROQ, coreROQ and ZnF show a single complex band with *FARSA*-AU RNA in EMSAs (**Figure 6A** and **Supplementary Figure 8A**). The ROQ and ZnF domains bind *FARSA*-AU with nanomolar and micromolar affinities, respectively; thus confirming earlier findings for ROQ(27) and the NMR and ITC data obtained in this study for the ZnF. EMSAs with RNA mutant versions (**Figure 6A** and **Supplementary Figure 8A**) confirm binding sites of the two domains: Mutating either the stem-loop or the AU-stretch abolishes complex formation of the ROQ domain or ZnF, respectively (**Supplementary Figure 8A**). Hence, the EMSAs unambiguously confirm that the ROQ domain interacts with the stem-loop structure(4,27) and that the ZnF binds the unstructured AU-rich stretch of *FARSA*-AU. The tandem domain constructs show transitions at approx. 200 and 400 nM for extROQ-ZnF and coreROQ-GS-ZnF, respectively, which indicates that the ROQ domain is the driving force for complex formation. EMSAs with mutant RNAs and tandem domain Roquin proteins showed transitions corresponding to either of the domains binding its designated target motif according to the shift pattern (**Figure 6A** and **Supplementary Figure 8A**). Conservation of individual affinities in the tandem domain context confirms that RNA-binding of ROQ and ZnF happens in an independent manner. The latter theoretically allows binding of two Roquin proteins to the same RNA molecule. To determine the stoichiometry of the complex, we monitored gel band shifts at high ligand concentrations. As a result, EMSAs of extROQ-ZnF at 5 µM RNA concentrations (approx. 20-fold > K_D_*(ROQ)*) suggested 1:1 binding in the complex (**Supplementary Figure 8C**), in line with our observations for the isolated ZnF in ITC (**Figure 2G**). Independent, but concerted binding of ROQ and ZnF domains to one RNA could provide higher target specificity.

**Figure 6.**
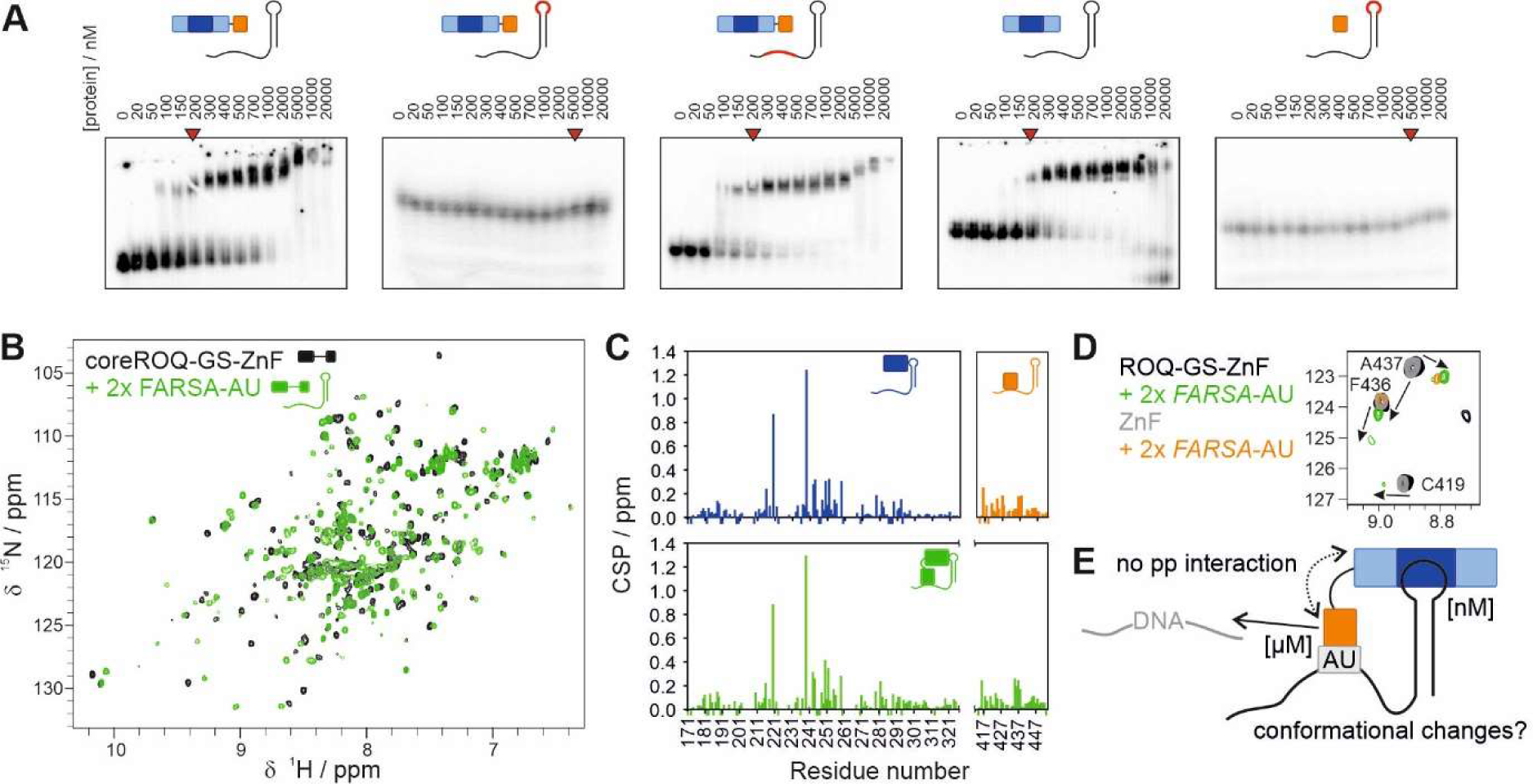
Natural target binding by ROQ domain and ZnF. **A)** EMSAs of single and tandem domain Roquin proteins with *FARSA*-AU wildtype and mutant RNAs. Red RNA cartoons depict mutant regions (see Supplementary Figure 8A). Red arrows indicate binding events. **B)** ^1^H,^15^N-HSQC spectra of apo coreROQ-GS-ZnF (black) and in presence of *FARSA*-AU RNA (green). **C)** CSPs of coreROQ, ZnF and the fusion construct (coreROQ-GS-ZnF) shown in B) upon titration of *FARSA*-AU RNA. **D)** Zoom-in of ^1^H,^15^N-HSQC spectra of apo coreROQ-GS-ZnF (black) and ZnF (grey) and in presence of *FARSA*-AU RNA (green and orange, respectively). Full spectra are shown in **Supplementary Figure 8B. E)** Model of bi-modal *FARSA*-AU recognition by ROQ and ZnF domains.

To corroborate the finding, we sought to obtain single-residue information on RNA binding in the two-domain context. We recorded NMR titration series of the isolated coreROQ domain, the ZnF and the tandem coreROQ-GS-ZnF construct with *FARSA*-AU RNA (**Figure 6B** and **Supplementary Figure 8B**). The ^1^H,^15^N correlation spectra reveal characteristic CSPs for the ROQ domain bound to a CDE (**Figure 6C**), known from previous studies(20), i.e. ROQ consequently interacts with *FARSA*-AU through the conserved binding mode. Similarly, the CSPs of the ZnF agree well with those we observed for the UAUUAUAU RNA (compare **Figure 2D** and **Figure 6C**), confirming the conserved binding mode for AU-rich RNA. In the context of the two-domain construct the CSP pattern is retained, from which we conclude that binding modes are maintained and the two domains do not modulate each other in RNA binding. While the isolated ZnF binds to RNA in an almost pure fast-exchange regime, the majority of peaks in coreROQ-GS-ZnF is severely line-broadened for both domains at substoichiometric RNA amounts. This line-broadening can be partially rescued when RNA is added in excess. Binding therefore occurs in the fast-to-intermediate exchange regime, in-line with nanomolar to low micromolar binding. The ROQ domain, thus dominates complex formation in the two-domain context. Since the exchange regime of the ZnF is also changed, we speculate that engagement of the ZnF with RNA in presence of the ROQ domain is entropically/statistically favoured, while the short linker (7 aa) keeps the domains independent. This is also supported by larger CSPs for ZnF residues in the coreROQ-GS-ZnF context (**Figure 6D**). Yet, we wondered whether the protein linker imposes conformational restrictions on the RNA site. Therefore, we compared *FARSA*-AU gel shifts of individual ZnF and extROQ domains with the extROQ-ZnF fusion construct (**Supplementary Figure 8D**). A mixture of both domains led to the same shift as the extROQ domain only, which is likely caused by a neglectable impact on the hydrodynamic radius by the small ZnF compared to the large ROQ domain (**Supplementary Table 1**). However, the tandem domains cause a larger shift than extROQ only. This suggests that simultaneous binding of ROQ and ZnF increases the hydrodynamic radius of the complex. Our NMR and EMSA data support a model where both ROQ and ZnF recognize their target sequences in *FARSA*-AU (**Figure 6E**). While binding modes are independent, ROQ facilitates ZnF:RNA interaction.

## DISCUSSION

RNA-binding proteins often utilize multiple RNA-binding domains to increase affinity or specificity for their RNA targets, integrating sequence and shape recognition(23). The ROQ domain of Roquin-1 recognizes stem-looped structures, which act as mRNA decay signals (4,5). The sheer number of decay elements in the genome(6) implies that co-existing *cis*-elements in mRNAs contribute to selectivity by Roquin. Those could be nearby dsRNA regions engaged by the dedicated extended ROQ domain(21,27). More likely, the Roquin ZnF contributes to the recognition of a combined motif or aids in motif engagement for the ROQ domain. Experimental evidence for the latter has been missing, mainly due to the lack of high-resolution information and the systematic definition of a consensus target sequence of the isolated ZnF. We have addressed both determinants based on the solution structure of the ZnF and its RBNS-derived precise motif, subsequently refined by NMR.

Similarly to ROQ, the Roquin ZnF is ultraconserved among vertebrates (**Supplementary Figure 1A**), but it is lacking in *C. elegans*, which indicates a specific role in the development of highly evolved adaptive immune systems. Its canonical CCCH zinc cluster, found in ZnF domains of other immune-relevant ZnF proteins(59), is C-terminally extended by a 12-residue helix, which has escaped earlier protein (secondary) structure predictions. We assume this helix is crucial for the fold and stability of the Roquin ZnF and inevitable for a protein construct accessible to *in vitro* and structural studies. Unexpectedly, the helical part seems completely devoid of RNA interactions (*see also below*). However, we do not rule out additional RNA-interactions in a larger functional context as shown for other ZnFs(60).

We were not able to provide a complex structure of the Roquin ZnF with RNA. However, the enrichment of a 6-mer in RBNS implies an extended interface with the ZnF, which involves the entire bottom part, but does not per se include the helical extension. Based on findings in other ZnF-RNA high-resolution structures we suggest the highly conserved aromatic residues Y414, Y417 and F436 undergo major conformational rearrangements during interaction with the AU-rich motif, which lead to a base-stacking pattern supported by charge-charge interactions (through K415). Similar types of interactions have e.g. been seen in the Tis11d-RNA complex as well as for Regnase-4 ZnF interacting with RNA(61). Such a scenario is in line with the observed CSP patterns and also supported by the observed differential CSP trajectories and magnitudes when comparing RNA and DNA (**Supplementary Figure 3C and D**). In fact, this type of complex formation with RNA has been discussed as an evolutionarily adapted feature of CCCH-type ZnFs(61). A structural alignment of the Roquin-1 ZnF with ZnF-RNA complexes of Regnase and Tis11d supports the assumption of an overall similar interaction with RNA(49,61), excluding the involvement of the helical extension (**Supplementary Figure 1E**).

Single ZnF domains, like exemplified here by Roquin-1, provide a comparably small surface for interaction with ssRNAs, while the observed interface approximately scales with the number of ZnFs involved(49,62). This fact is also reflected in the number of engaging nucleotides. MBNL1 binds 3-mer RNA motifs(50), both FUS and Tis11d ZnFs were shown to recognize 4-mers(49). The latter one is a CCCH-type ZnF and similarly binds AU-rich elements, while e.g. the Nab2 ZnF is able to interact with a stretch of purines(62). Interestingly, the Tis11d tandem ZnFs bind a UAUU motif each(49), reminiscent of our findings of single adenosines embedded in a poly(U) context (**Figure 4**).

While the ROQ domain binds with (low) nanomolar affinity(4), the ZnF affinity for an RBNS-derived RNA (**Figure 2**) is 14 µM. This represents a typical affinity for single RNA-ZnF complexes(23,51), and is supported by our NMR titration data. Nanomolar affinities can be achieved by two or more ZnFs binding to one RNA(50,62). Copies of (different) ZnFs in one protein increase avidity and specificity or serve as a molecular ruler in spacing other RNA-binding domains. Further, motif length can affect the apparent affinity and, in fact, an avidity effect could enhance binding of the Roquin ZnF to natural targets, where AREs are usually repetitions, e.g. of our tested motif(63). The moderate affinity of single ZnFs is required for target search(64,65), where high affinities impede sliding across multiple registers(66),(64). Similarly, ARE-binding ZnFs, could enhance target search and affinity, i.e. by dynamic binding to AU-rich RNA adjacent to CDEs or ADEs. We observed dynamic binding of the Roquin ZnF to poly(U) stretches in our NMR experiments (**Figure 5**)(57,58). Of note, no systematic study has analyzed the co-appearance of stem-loop elements with AREs in mRNA 3’UTRs. Instead, we currently rely on single case studies(2). However, the lack of AREs in Roquin targets like *Ox40*, *Nfκbid* or *Tnfα* points at a context- and target-dependent function of the ZnF.

Originally described as transcription factors(67), meanwhile, ZnFs binding to dsRNA or RNA-DNA hybrids have been discovered(68). We found that the Roquin ZnF can bind both RNA and DNA, in line with a prior study(56). Interestingly, our NMR analysis reveals that the ZnF binds both types of nucleic acid through the same interface, albeit with different modes of interactions (**Figure 3**). These differences point at specific recognition of 2’-OH groups and discrimination between uracils and thymines. While dual DNA- and RNA-binding by proteins is a recurring feature(69), e.g. for transcription factors(70), it usually requires multiple domains(54,71) or alternatively spliced protein variants, and rarely exploits a partially overlapping interface(72). Due to its cytoplasmic localization, a DNA-binding role of Roquin is speculative, and its DNA-binding capacity might be an evolutionary remnant. Nonetheless, we find a systematic parallelism in DNA- and RNA-binding regarding the tested motifs and thus decided to use DNA as a fast and cost-efficient surrogate to refine the RBNS-derived RNA lead motif via a PCA by NMR (**Figure 4**) (73). It allowed us to systematically reconstruct an RBNS-derived motif and to evaluate individual positions for affine ZnF binding. This approach can be easily applied to other cases of RBD-RNA motif refinement by (cost-)efficient and fast, but highly resolved structural biochemistry.

We showed that the Roquin ZnF interacts with AU-rich RNA sequences adjacent to a CDE element in presence of the ROQ domain (**Figure 6E**). Our data show that both domains engage with unique RNA interfaces, as observed in our RNA mutants, and the binding mode is preserved in presence of both protein domains. While its micromolar affinity suggests a subordinated role in steering towards a combined canonical target motif together with the ROQ domain, it may increase specificity for a subset of Roquin targets. Indeed, a too affine ZnF-ssRNA interaction will counteract the functionally relevant interaction of ROQ with stem-loop elements, thereby limiting a guiding function. However, our data do not rule out Roquin-mediated mRNA regulation solely based on ZnF-RNA interactions, e.g. through binding to an expanded ARE and in the absence of CDEs and ADEs.

In a different scenario, individual domains of a MD-RBP form combined RNA-binding interfaces through domain-domain interactions per se(74). For Roquin, a recent crystal structure suggested an interaction between ROQ and the RING domain(22). An interaction of ROQ and the ZnF has not been described because of the lack of structural information of the ZnF, but was discussed in a functional context (7,75). Our successful production of the ZnF alone and in a fusion construct with ROQ now allowed to address this question. We did not observe domain-domain interactions between ROQ and ZnF domains, which could have affected the combined target recognition. This may indirectly support previous crystallization attempts, where in a three-domain construct (RING-ROQ-ZnF) no electron density was observed for the ZnF(22). We conclude that ROQ and ZnF are structurally uncoupled, yet they are close in space, connected by a short (7 aa) linker. Structural independence of the ZnF has been observed for other MD-RBPs before(23,76). The short linker potentially requires conformational rearrangements of the RNA for complex formation with both protein domains, as we conclude from our gel shift assay (**Supplementary Figure 8D**)(77,78), which indicate a possible compaction of the RNP as it was also observed in Regnase RNPs(61).

Finally, we need to consider a role for the Roquin ZnF in mediating intermolecular protein-protein interactions (PPI), e.g. through the newly identified helical extension. Indeed, PPIs have prior been found mediated through CCHC or CCHH type ZnF(11,12). For Roquin proteins, it remains to unambiguously define their oligomeric state *in vivo* (7). As such, the final relevance of the ZnF domain could potentially rise in case of dimeric target recognition. A respective RNA motif may therein mediate ZnF dimerization, while our data do not suggest ZnF oligomers per se, as e.g. seen by NMR relaxation (not shown). The helical extension may also serve as platform for other proteins. A recent study has revealed the crucial interaction of Roquin with the functionally related Regnase (79), where the two proteins may use their similarly typed ZnFs for additional interactions or for a combined initial engagement with an RNA target. Finally, while our group has shown that both RING and ZnF do not interact with ROQ in solution, we do not exclude intramolecular interactions of the ZnF helix with other parts of the full-length protein.

Altogether, the herein determined findings essentially complement the structural information on the RNA-interacting Roquin N-terminal multi-domain region and its combined target consensus RNA. The data suggest how Roquin integrates RNA shape- and sequence features for a fine-tuned RNA-specificity. Our work also serves as a technical template for exploiting the combined strength of RBNS and NMR, followed by RNA-protein biochemistry, applicable to any RBD-RNA question.

## ACCESSION NUMBERS

NMR chemical shifts of Roquin ZnF have been deposited at the BioMagResBank (BMRB) under ID 52226. The structural ensemble is deposited in the Protein Data Bank (PDB) under entry ID 8RHS. RBNS NGS data have been deposited with the Gene Expression Omnibus database (GEO; www.ncbi.nlm.nih.gov/geo/) and are available under the accession number GSE253536. Token for reviewer access is abcposaerpojbed.

## SUPPLEMENTARY DATA

Supplementary Data are available at XXX online.

## ACKNOWLEDGEMENT

We acknowledge excellent technical support by Katharina Targaczewski and thank Anna Wacker for support with TOCSY measurements. We thank the Sattler lab and the Bavarian NMR center for the opportunity to record some of the initial NMR experiments of this study. Simon Schäfer is to be acknowledged for his support in cloning of ROQ-ZnF fusion constructs.

## SUPPLEMENTARY MATERIAL

### SUPPLEMENTARY TABLES

**Supplementary Table 1:** Roquin protein constructs used in this study, their boundaries and molecular weights.

| Protein | Boundaries | Number of amino acids | MW / kDa |
| --- | --- | --- | --- |
| ZnF | 411-454 | 44 | 5.2 |
| coreROQ | 171-326 | 156 | 17.8 |
| extROQ | 89-404 | 316 | 35.4 |
| coreROQ-GS-ZnF | 171-326-GGGGS-411-454 | 205 | 23.3 |
| extROQ-ZnF | 89-454 | 366 | 41.3 |

**Supplementary Table 2:**
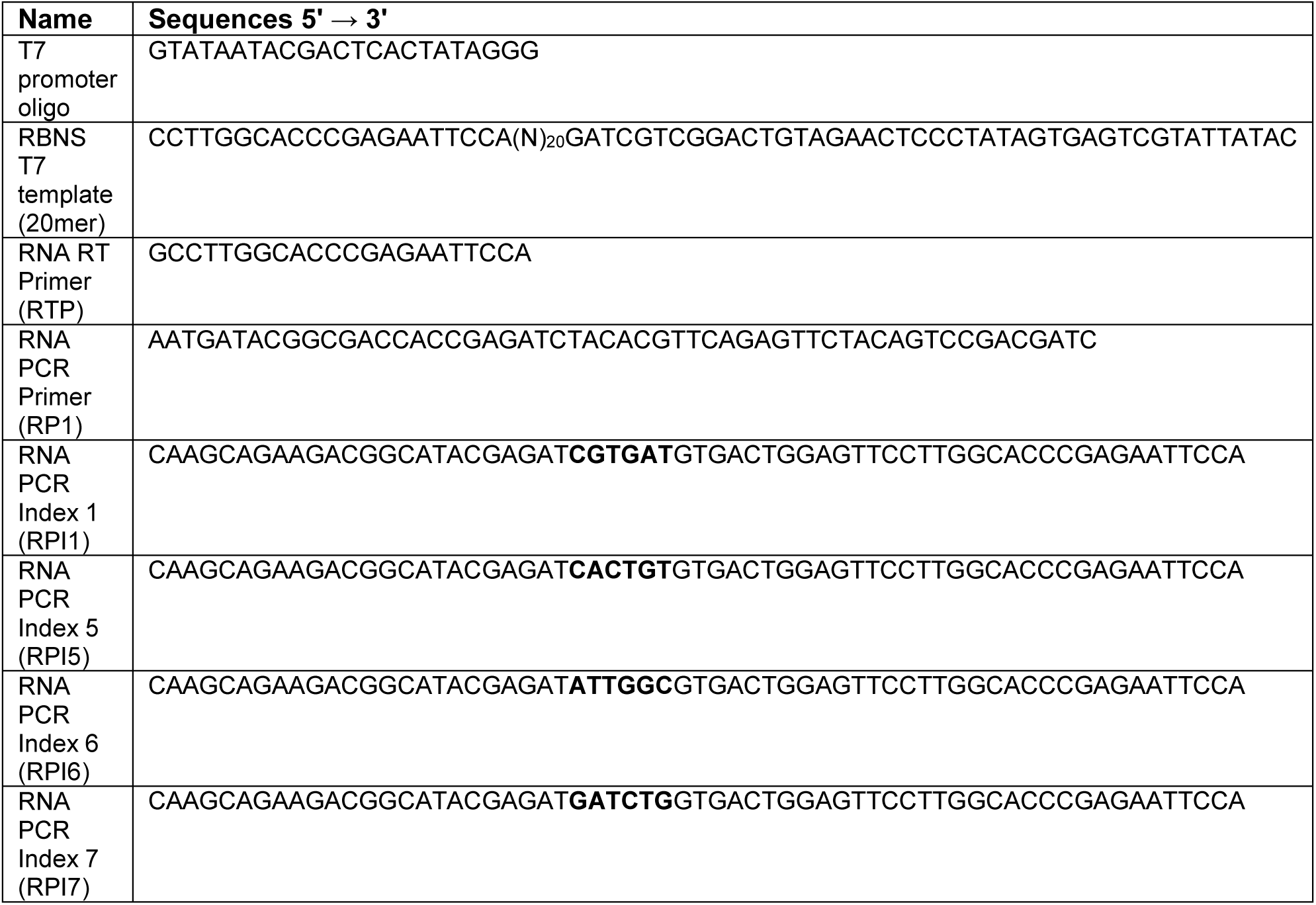
Oligonucleotides used for the RBNS assay.

**Supplementary Table 3:** Indices used for illumina sequencing.

| Sample Name | i7 Index Name | i7 index Sequence |
| --- | --- | --- |
| input | RPI1 | ATCACG |
| ZnF-1-uM | RPI5 | ACAGTG |
| ZnF-5-uM | RPI6 | GCCAAT |
| ZnF-20-uM | RPI7 | CAGATC |

**Supplementary Table 4:**
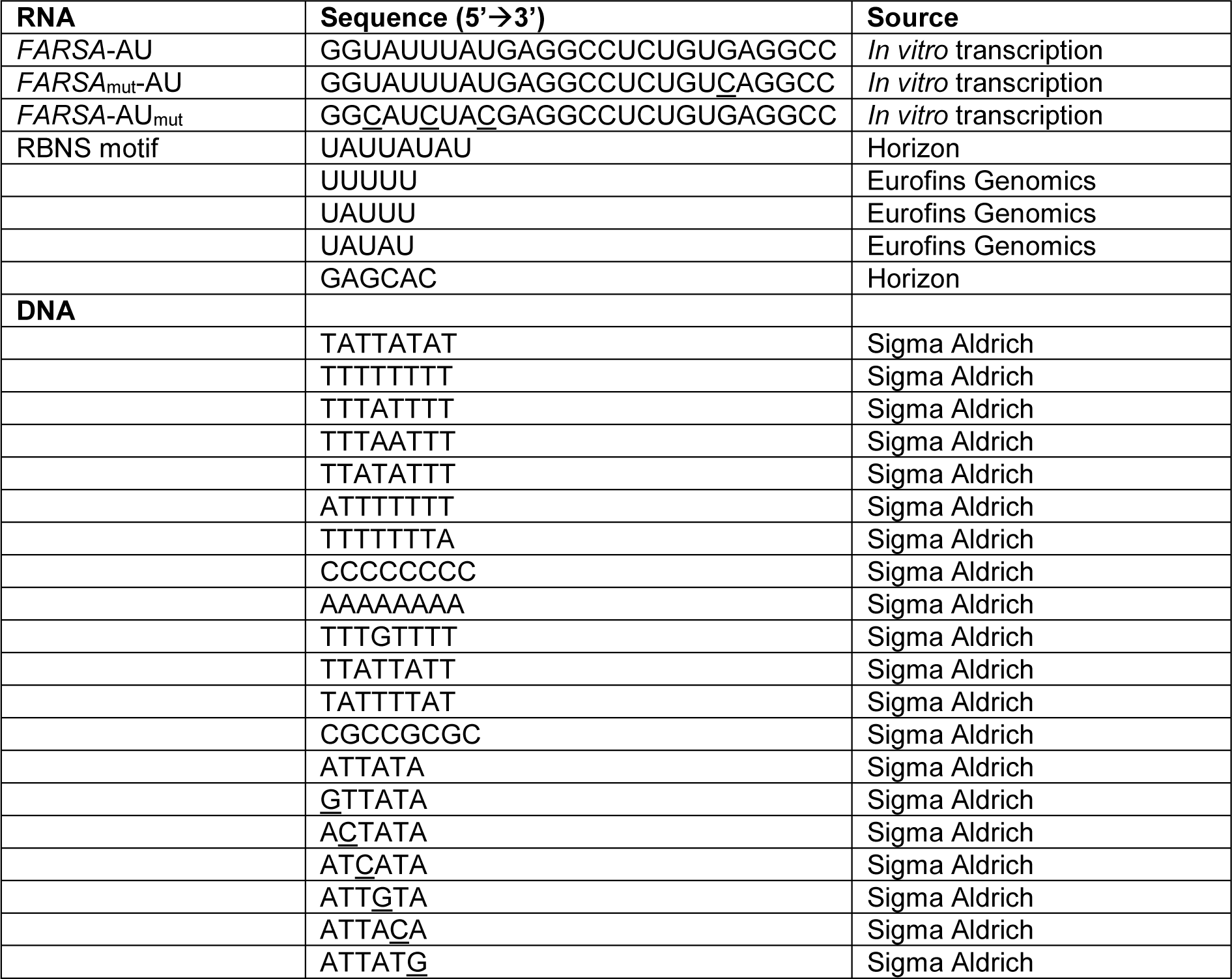
RNAs and DNAs used in this study. Underlined residues are mutated with respect to reference sequence.

**Supplementary Table 5:** Provided as separate file.

### SUPPLEMENTARY FIGURES

**Supplementary Figure 1.**
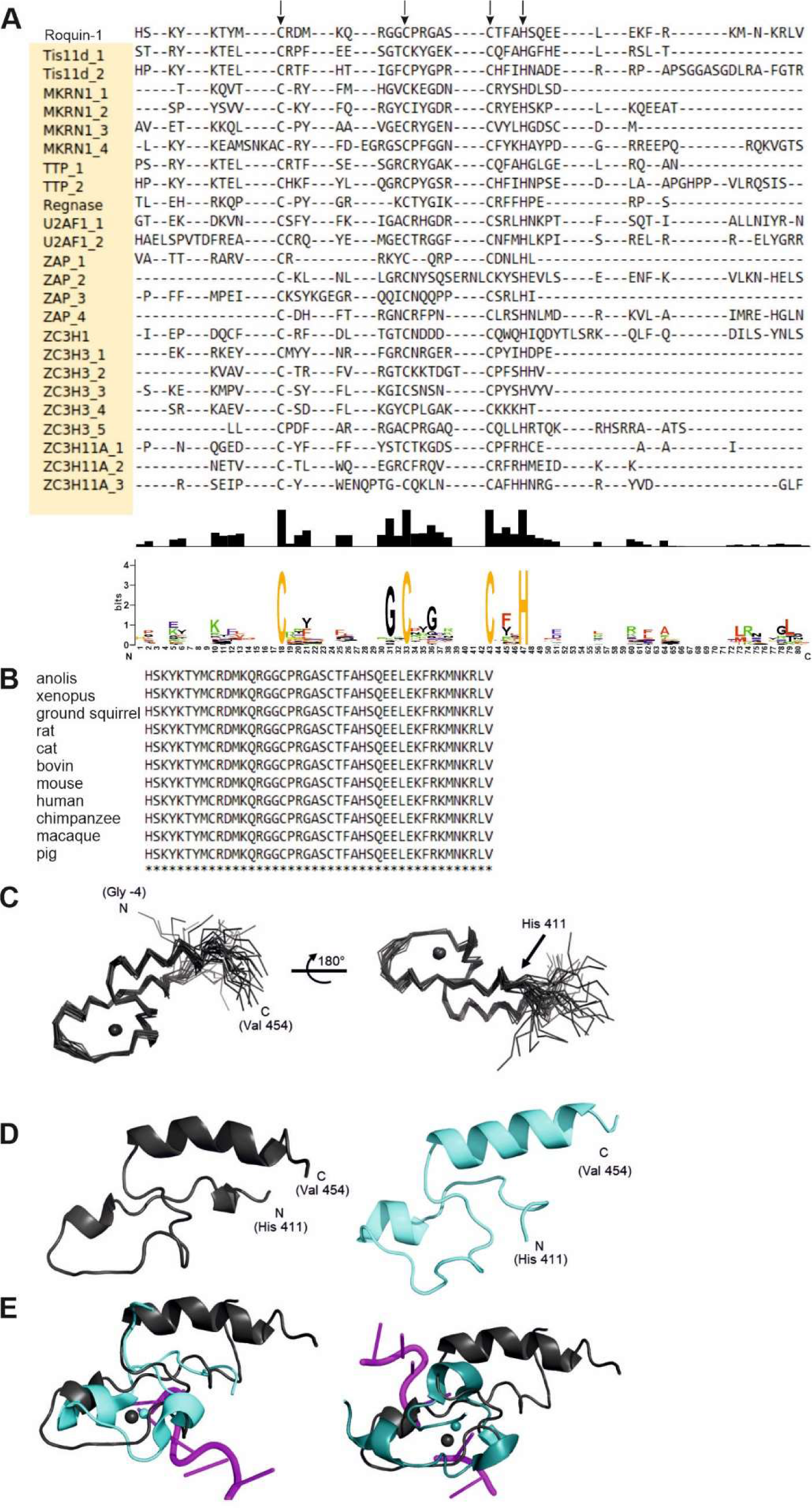
Sequence conservation of the Roquin-1 ZnF and its structure. A) Alignment of Roquin ZnF with other human CCCH-type ZnFs. Given ZnFs were aligned to residues 411-454 of Roquin ZnF. Arrows indicate anchor points used for alignment. Shown below are the conservation score and the consensus motif derived from the alignment. **B)** Sequence alignment of Roquin-1 ZnF from multiple species. **C)** Structure of the human Roquin-1 ZnF domain. Ribbon presentation of the 20 lowest-energy models shown in main text Figure 1. For orientation, the zinc ion is shown as sphere as derived from structure determination when used as pseudo atom. The natural sequence start (His 411) is shown in the right view, while the full construct is depicted including the TEV-cleavage derived (-4) - GAMA - (-1) overhang. **D)** Comparison of the lowest energy representative including residues 411-454 with a structural model of the Roquin ZnF as taken from AlphaFold (48) entry Q5TC82 (human Roquin-1, showing residues 411-454). **E)** Alignment of apo Roquin-1 ZnF (black) with ZnFs of Tis11d (PDB: 1RGO (49); left, cyan) and Regnase (PDB: 7NDJ (61); right, dark cyan), both in RNA-bound form. The RNA is shown in magenta.

**Supplementary Figure 2.**
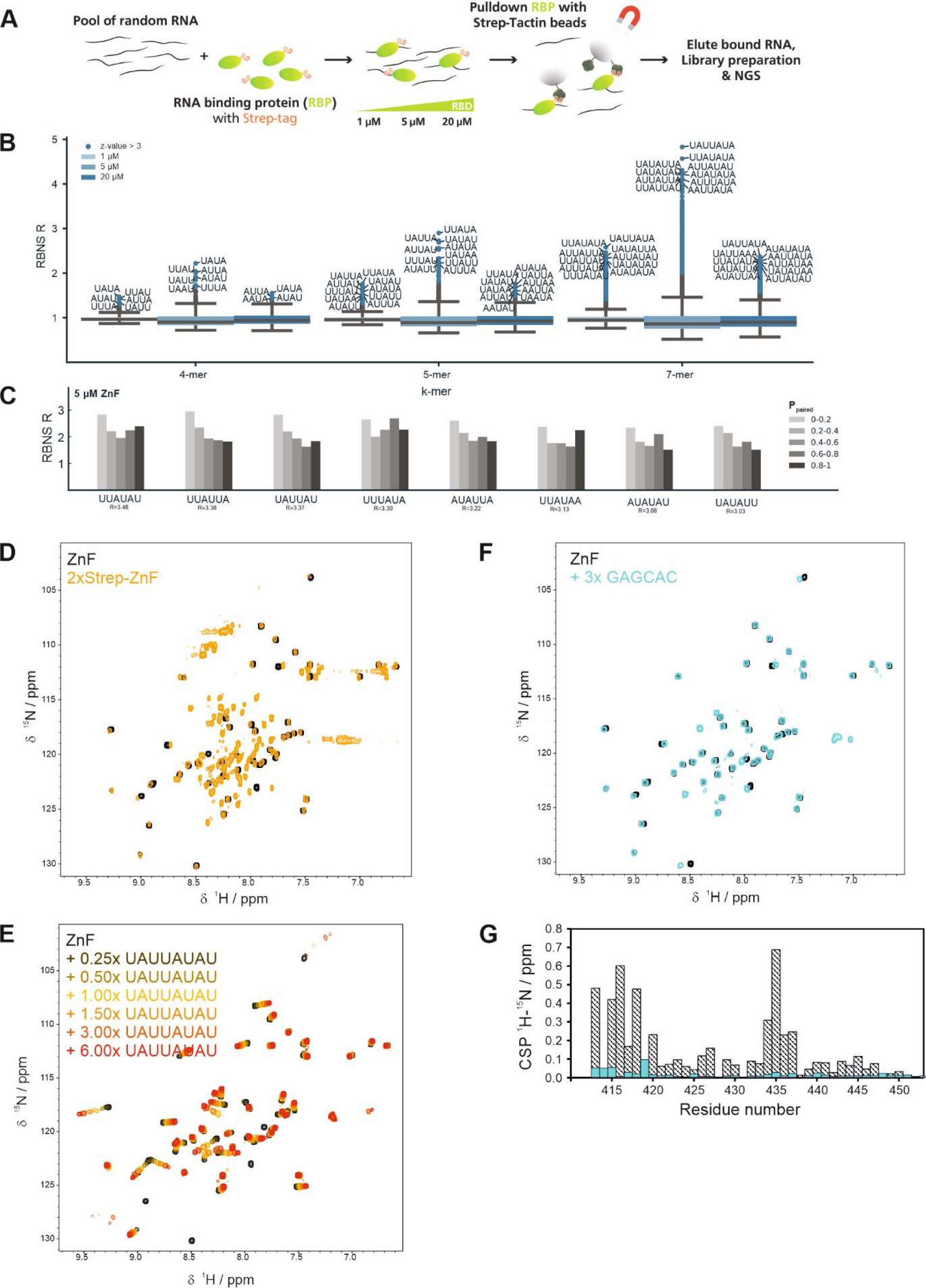
AU-RNA binding of Roquin ZnF. **A)** Experimental design to identify a binding motif for the ZnF domain by RBNS. **B)** Enrichment of different k-mers (k = 4,5,7) at 1, 5 and 20 µM ZnF concentration. Values greater than three standard deviations above the mean are highlighted. The ten most enriched motif sequences are given. **C)** Structural features of the 3^rd^ to 10^th^ most enriched 6-mer motifs at 5 μM ZnF. D) ^1^H,^15^N-HSQC spectra of ZnF (black) and a Twin-Strep-tagged ZnF construct (yellow). **E)** ^1^H,^15^N-HSQC spectra of apo ZnF (black) and in complex with increasing concentrations of AU-rich RNA (yellow to red). **F)** ^1^H,^15^N-HSQC spectra of apo ZnF (black) and in the presence of control RNA. **G)** CSPs of F) shown in turquoise compared to CSPs of a 1:3 complex of AU-rich RNA (shaded black) from **E)**.

**Supplementary Figure 3.**
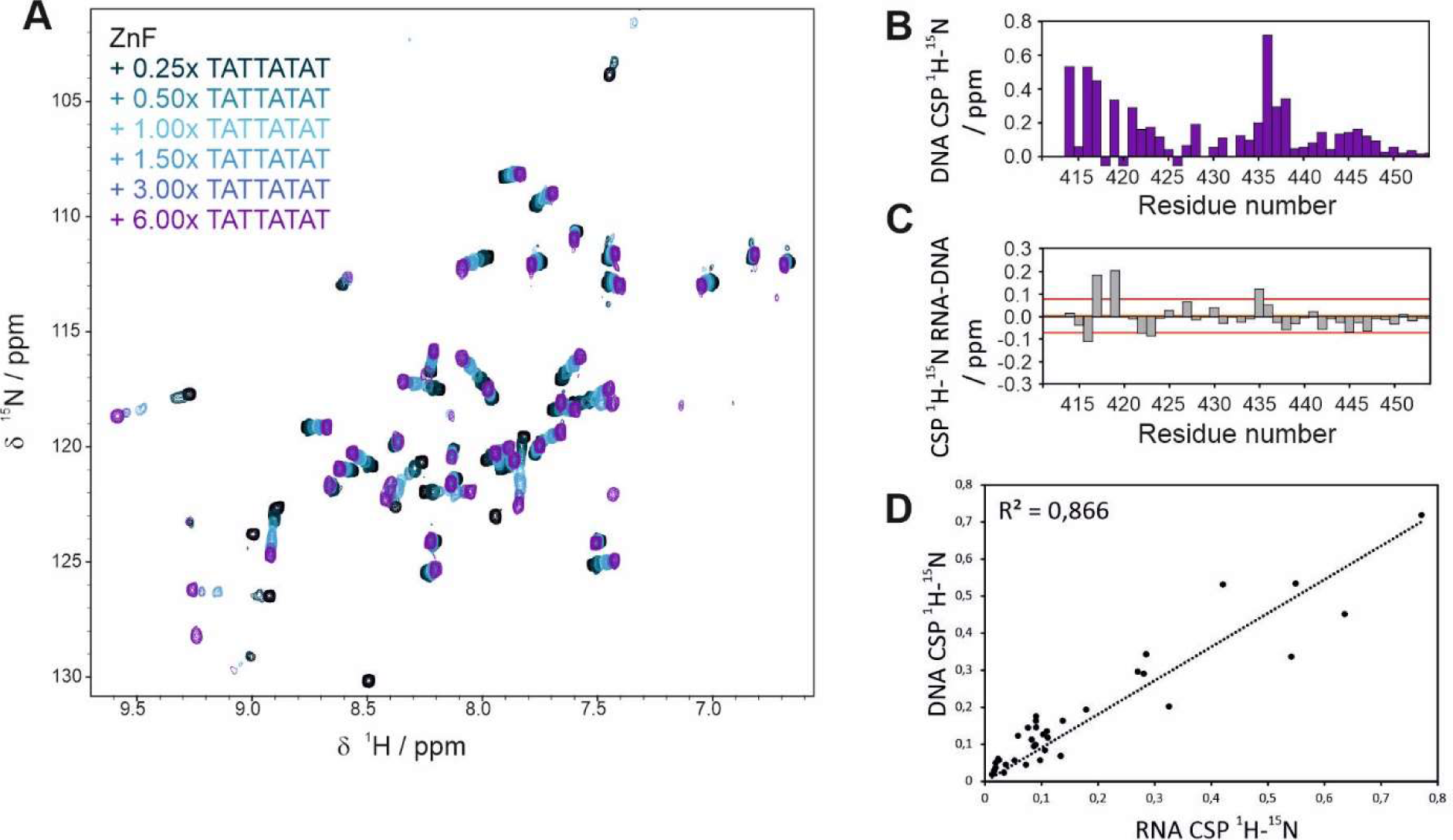
Key ZnF residues involved in RNA and DNA binding. **A)** ^1^H,^15^N-HSQC spectrum of apo ZnF (black), and in presence of increasing DNA concentrations (blue to purple). **B)** CSPs of titration shown in **A)**. Negative values indicate non-trackable residues. **C)** CSP difference plot of RNA and DNA. Red lines indicate average CSPs +/- one standard deviation. **D)** Correlation plot of DNA and RNA CSPs.

**Supplementary Figure 4.**
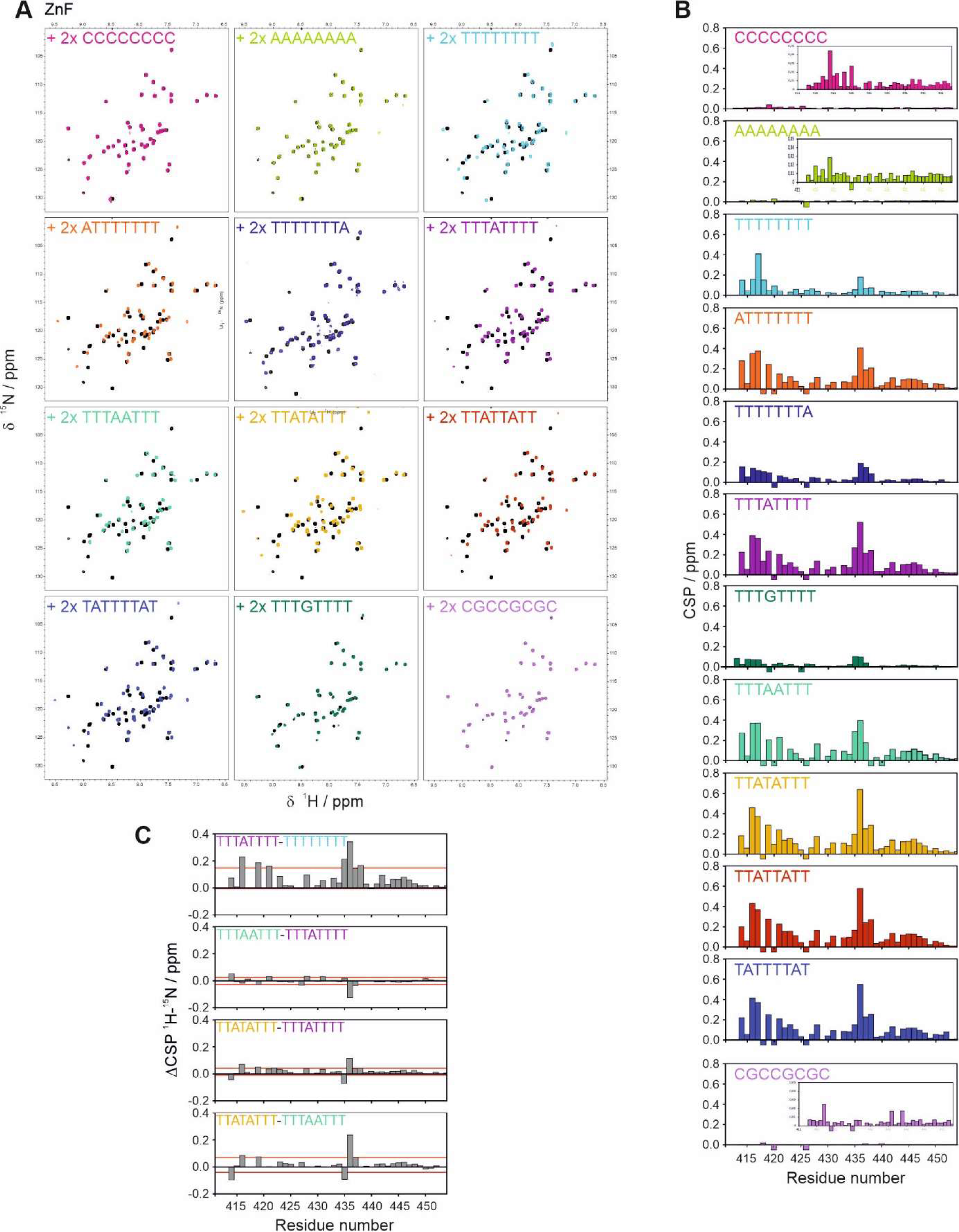
Sequence requirements for nucleic acid binding by Roquin ZnF. **A)** ^1^H,^15^N-HSQC spectra of apo ZnF (black) and in complex with DNA oligos (colored). **B)** CSPs of complexes shown in A). **C)** CSP difference plots of selected DNA titrations shown in **A)** and **B)**. Red lines indicate average CSPs +/- one standard deviation.

**Supplementary Figure 5.**
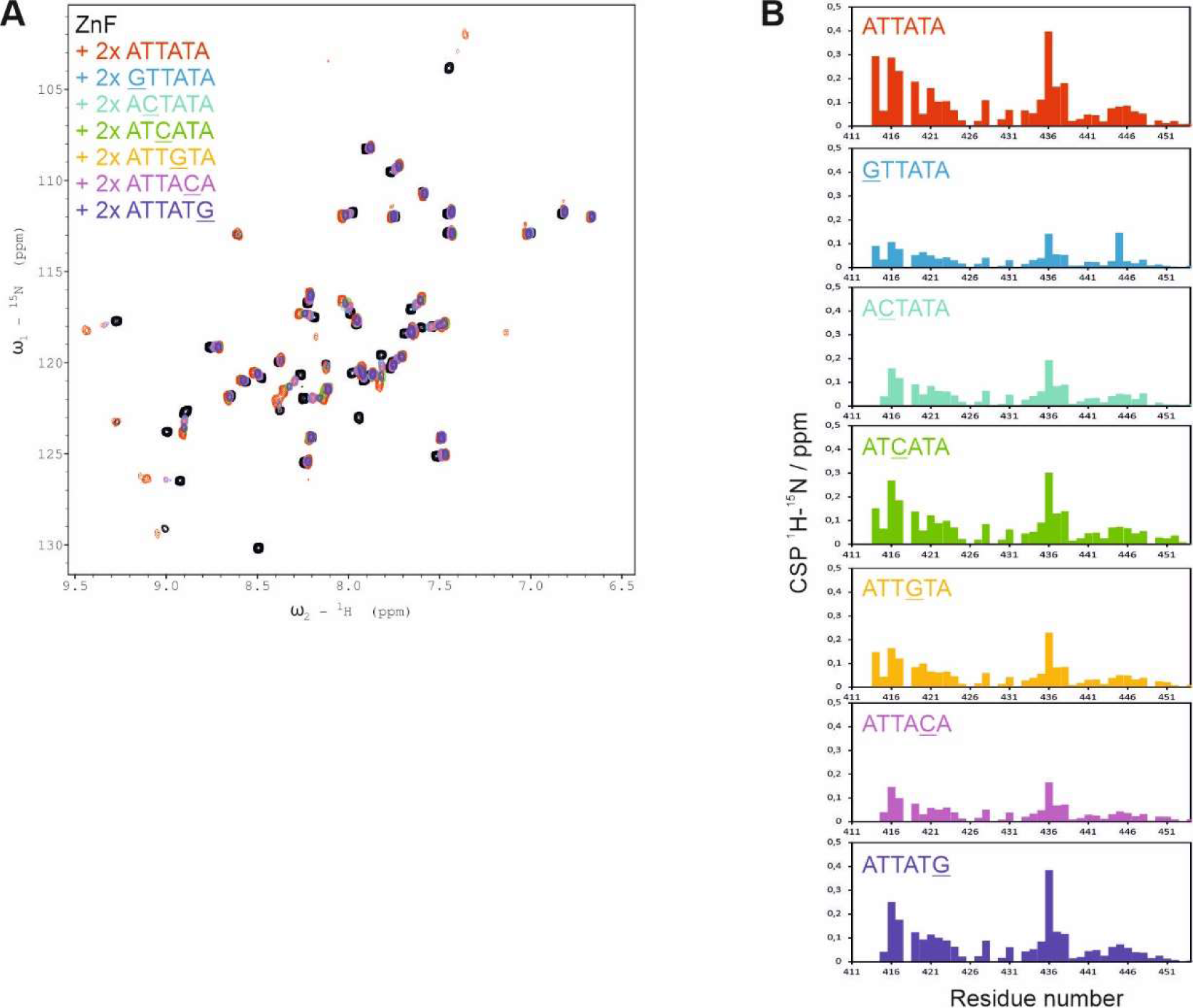
Positional hierarchy in the RBNS motif. **A)** ^1^H,^15^N-HSQC spectra of apo ZnF (black) and in complex with DNA variants of RBNS motif (colored). Underlined residues indicate mutation site with respect to RBNS motif. **B)** CSPs of DNA titrations shown in **A)**.

**Supplementary Figure 6.**
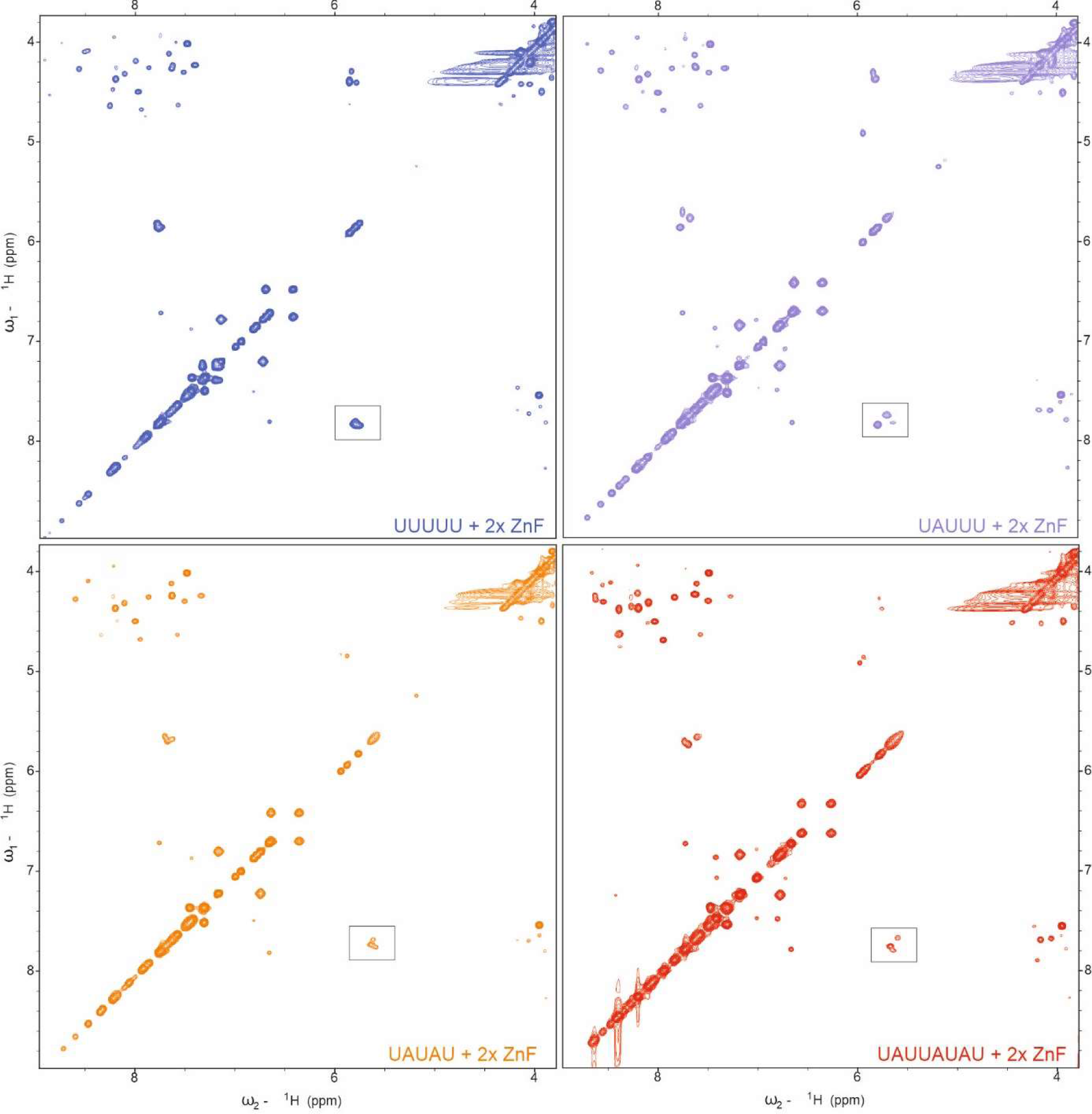
Multiple register binding in poly(U) RNAs. ^1^H,^1^H-TOCSY spectra of RNAs complexed with ZnF. Boxes indicate zoom-ins shown in Figure 5A.

**Supplementary Figure 7.**
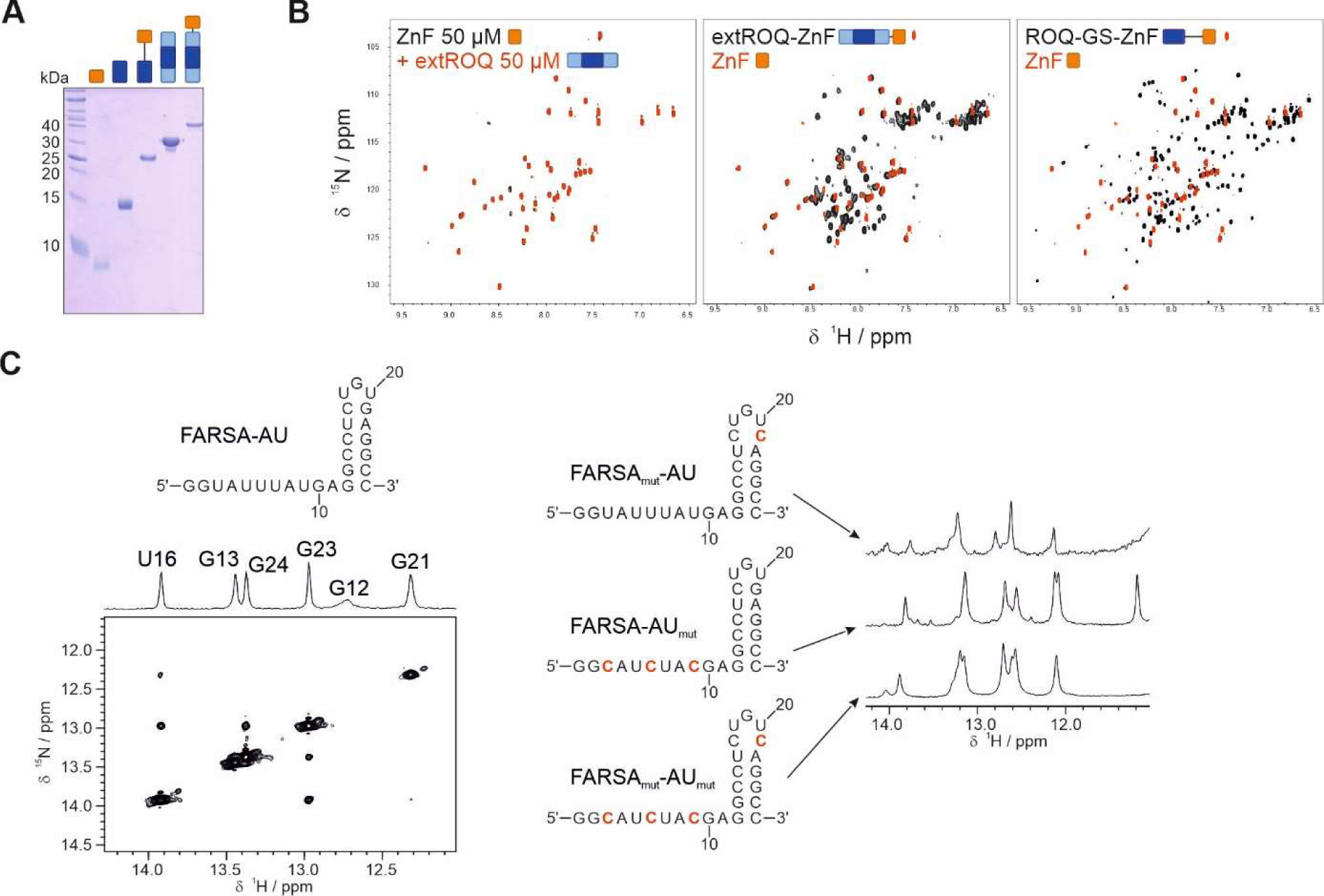
Protein domain independence and secondary structure of a Roquin target *cis*-element. **A)** SDS-PAGE of protein constructs used in this study (**Supplementary Table 1**). **B)** ^1^H,^15^N-HSQC spectra of apo ZnF (black) and in presence of extROQ (red; left), and extROQ-ZnF and coreROQ-GS-ZnF (black; center and right, respectively) overlayed with ZnF (red). **A)** ^1^H,^1^H-NOESY of imino region of wildtype *FARSA*-AU RNA (left) and derived secondary structure. Mutant versions of *FARSA*-AU RNAs are shown on the right with corresponding imino ^1^H spectra. Mutant sites are highlighted in red.

**Supplementary Figure 8.**
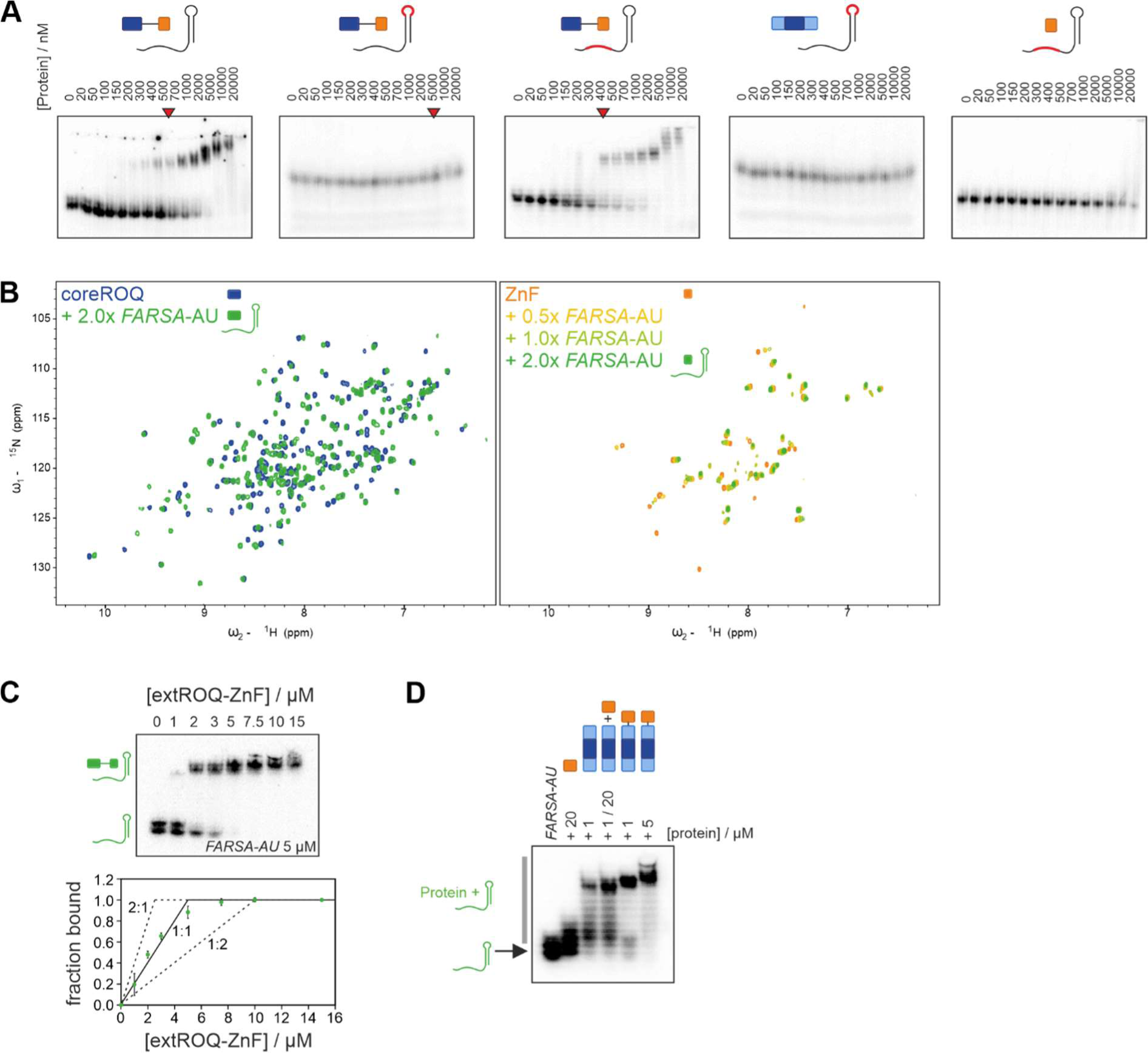
Natural target binding by ROQ and ZnF domains. **A)** EMSAs of single and tandem domain Roquin proteins with *FARSA*-AU wild type and mutant RNAs. Red RNA cartoons depict mutant regions. Red arrows indicate binding events. **B)** ^1^H,^15^N-HSQC spectra of apo coreROQ (blue; left) and ZnF (orange; right) and in presence of *FARSA*-AU (green and yellow, respectively). Corresponding CSPs are shown in Figure 6 **C**. **C)** Representative stoichiometric EMSA of extROQ-ZnF vs. *FARSA*-AU at 5 µM RNA. Below fractions bound from triplicates are plotted together with theoretical curves for 2:1, 1:1 and 1:2 binding. **D)** EMSA of *FARSA*-AU with single, tandem or mixed Roquin domains. Cartoons depict proteins used in each lane. The arrow marks free RNA while the grey bar labels RNPs.

